# Network analysis allows to unravel breast cancer molecular features and to identify novel targets

**DOI:** 10.1101/570051

**Authors:** Aurora Savino, Lidia Avalle, Emanuele Monteleone, Irene Miglio, Alberto Griffa, Giulia Accetta, Paolo Provero, Valeria Poli

**Affiliations:** Department of Molecular Biotechnology and Health Sciences, Molecular Biotechnology Center, University of Turin, Via Nizza 52, 10126 Turin, Italy; Center for Translational Genomics and Bioinformatics, San Raffaele Scientific Institute, 20132 Milan, Italy

## Abstract

The behaviour of complex biological systems is determined by the orchestrated activity of many components interacting with each other, and can be investigated by networks. In particular, gene co-expression networks have been widely used in the past years thanks to the increasing availability of huge gene expression databases. Breast cancer is a heterogeneous disease usually classified either according to immunohistochemical features or by expression profiling, which identifies the 5 subtypes luminal A, luminal B, basal-like, HER2-positive and normal-like. Basal-like tumours are the most aggressive subtype, for which so far no targeted therapy is available.

Making use of the WGCNA clustering method to reconstruct breast cancer transcriptional networks from the METABRIC breast cancer dataset, we developed a platform to address specific questions related to breast cancer biology. In particular, we obtained gene modules significantly correlated with survival and age of onset, useful to understand how molecular features and gene expression patterns are organized in breast cancer. We next generated subtype-specific gene networks and in particular identified two modules that are significantly more connected in basal-like breast cancer with respect to all other subtypes, suggesting relevant biological functions. We demonstrate that network centrality (kWithin) is a suitable measure to identify relevant genes, since we could show that it correlates with clinical features and that it provides a mean to select potential upstream regulators of a module with high reliability. Finally, we showed the feasibility of adding meaning to the networks by combining them with independently obtained data related to activated pathways.

In conclusion, our platform allows to identify groups of genes highly relevant in breast cancer and possibly amenable to drug targeting, due to their ability to regulate survival-related gene networks. This approach could be successfully extended to other BC subtypes, and to all tumor types for which enough expression data are available.

## Introduction

The behaviour of complex systems emerges from the orchestrated activity of many components interacting with each other. Thus, networks are a valuable construct to investigate the property of biological systems (Barabási & Oltvai, 2004). In this context, networks are typically defined by genes as nodes and regulatory relationships, actual or potential, as edges. Network inference can be based on high-throughput data only, in which case it is completely unbiased, or rely on prior information (Chasman, Siahpirani, & Roy, 2016). Both methods have advantages and downsides. The unbiased method allows the exploration of the whole gene network in a plethora of conditions, while introducing specific experimentally-driven regulatory interactions, despite significantly increasing network information content and reliability, is limited to a few nodes, edges and contexts for which experimental data are available (Chasman et al., 2016). As a consequence of the increasing number of available transcriptomic data, gene co-expression networks are amongst the most widely studied ones (Zhao et al., 2010). They are considered a proxy for gene regulatory networks, with the assumption that in most cases transcription factors and the genes they positively regulate are correlated at the transcriptional level. Nevertheless, there is no attempt to draw direct causal relationships among the genes in the network in the form of directed edges. Additionally, interactions in eukaryotes are difficult to infer, as relationships between the expression of regulators and targets are context-dependent due to the complex combinatorial nature of eukaryotic transcriptional regulation (Neph, Stergachis, Reynolds, Sandstrom, & Borenstein, 2012), suggesting that the most reliable network constructions can be obtained by separating the samples according to specific conditions/groups.

Networks have been used to understand disease mechanisms (Emilsson et al., 2008; Feldman, Rzhetsky, & Vitkup, 2008; Furlong, 2013; Goh, Cusick, Valle, Childs, & Vidal, 2007; Malod-dognin et al., 2019), and define therapeutic drugs and their targets (Isik, Baldow, Cannistraci, & Schroeder, 2015; L. Liu et al., 2018; Yamanishi, Araki, Gutteridge, Honda, & Kanehisa, 2008). In particular, since complex systems are very robust to component perturbations, network structure can inform on specific genes that are fundamental for network integrity (Albert, Jeong, & Barabási, 2000). In fact, the number of connections of a gene was shown to be correlated with its relevance in maintaining cell viability in *S. cerevisiae* (Jeong, Mason, Barabási, & Oltvai, 2001), and essential proteins have been found to be associated with hubs (i.e. highly connected genes) in human (Goh et al., 2007). This property is largely due to the scale-free topology of biological networks, meaning that most genes have low numbers of connections, while a few genes are highly connected (Albert et al., 2000).

Gene networks are likely to be organized in a modular fashion, with modules identifying groups of tightly topologically or functionally interconnected genes (Hartwell, Hopfield, Leibler, & Murray, 1999). Several methods have been proposed for modules inference (Saelens, Cannoodt, & Saeys, 2018), with clustering methods being particularly suited for global networks analyses and interpretation, despite not allowing the definition of finer grained modules with possibly overlapping genes. WGCNA (Weigthed Gene Coexpression Network Analysis) is a co-expression modules identification method based on clustering that assumes a scale-free topology and defines gene connectivity based on topological features (Langfelder & Horvath, 2008; B. Zhang & Horvath, 2005).

Breast cancer (BC) is one of the leading causes of mortality in women worldwide, for which no general efficacious treatment is available due to disease heterogeneity. BC is usually classified according to the immunohistochemical assessment of the expression of Progesterone and Estrogen Receptors (ER-positive BC), human epidermal growth factor receptor 2 (HER2-positive) and the proliferation marker Ki67 (Goldhirsch et al., 2009; Viale, 2012). Tumours negative for ER, PR and HER2 are defined as triple negative (TN). Alternatively, gene expression profiling of the tumour (Prediction Analysis of Microarray or PAM 50 assay) allows the classification in five molecular subtypes, partially overlapping with histological characterization: luminal A, luminal B, basal-like, HER2-positive and normal-like. These are correlated with prognosis and response to treatments (Perou et al., 2000). In particular, the basal-like (BL) subtype, which mostly correspond to TNBC, does not usually respond to targeted treatments such as hormonal blockage or Herceptin, and shows poor outcome and high frequency of TP53 mutations (Perou et al., 2000). Moreover, basal-like tumours show hyperactivation of the STAT3 and Wnt/PCP pathways. The TF STAT3 (Signal Tranducer and Activator of Transcription 3) is considered as an oncogene and correlates with poor prognosis. It is implicated in many functions fundamental for cancer progression such as proliferation, survival, invasiveness and metastasis (Avalle, Pensa, Regis, Novelli, & Poli, 2012; Yu, Lee, Herrmann, Buettner, & Jove, 2014). Survival of many basal-like cell lines was shown to require STAT3 activity (Muellner et al., 2015). Similarly, Wnt signalling has been implicated in malignant transformation (Logan & Nusse, 2004). Depending on the type of ligands involved, Wnt signalling has been classified as canonical or non-canonical, with the non-canonical Planar Cell Polarity (PCP) pathway being involved in defining cell shape and movement and inducing cancer progression (Klemm et al., 2011; Veeman, Axelrod, & Moon, 2003; Wang, 2009), particularly in basal-like BC.

Network inference has been applied to the study of breast cancer for the identification of prognostic modules and centrally connected genes as potential therapeutic targets (Clarke et al., 2013; Herschkowitz et al., 2010; X. Liu et al., 2007; Shi et al., 2017; Wolf, Lenburg, Yau, Boudreau, & Veer, 2014; Yang et al., 2014). Nevertheless, most published works are purely descriptive and do not exploit the wealth of information hidden in these networks for the definition and investigation of specific biological hypotheses. Additionally, the problem of differences in network structure across conditions has motivated the definition of differential co-expression analysis methods (Klein, Oualkacha, Lafond, & Bhatnagar, 2016; Yu, Zhao, Wang, Zhao, & Zhao, 2017; Zhu et al., 2017), but network structure and connectivity have never been compared across breast cancer subtypes. However, this investigation could drive the generation of new hypotheses about the molecular features conferring aggressiveness to the basal-like subtype.

Reconstructing breast cancer transcriptional networks, we developed a platform to address specific questions for BC in general and for specific BC subtypes: i) Which are the genes possibly driving clinical features? ii) Can we identify upstream regulators or downstream targets of specific signalling pathways? iii) Are there specific gene interactions explaining the phenotypic differences between cancer subtypes? Our platform represents a tool to address these and many other biological questions, identify key regulatory genes and drive experimentally-testable hypotheses.

## Results

### Network construction

We applied WGCNA (Weighted Gene Coexpression Network Analysis) (Langfelder & Horvath, 2008; B. Zhang & Horvath, 2005) to a wide primary breast tumour transcriptome dataset (METABRIC) (Curtis et al., 2012), defining 21 modules of co-regulated genes across all patients (Suppl. Fig. 1). Modules’ names were assigned by analysing the Hallmarks sets defined by the Molecular Signature Database (http://software.broadinstitute.org/gsea/msigdb), choosing the gene set displaying the most significant enrichment among the module’s genes. NC denotes the modules for which no significant enrichment was found. These groups of genes were confirmed to be robust, being significantly co-regulated both in an independent primary breast tumour dataset (TCGA-BRCA) and in a panel of breast cancer cell lines (http://cancergenome.nih.gov/, Daemen et al., 2013, Suppl. Fig. 2). Moreover, the relationships between genes within each module were confirmed by their enrichment in interacting proteins, as obtained from STRING protein interaction database (Jensen et al., 2009) (not shown). Based on these networks we developed two lines of investigation, aimed at identifying genes most likely involved in i) determining clinical features and ii) being part of selected signalling pathways linked to cancer aggressiveness.

**Figure 1.**
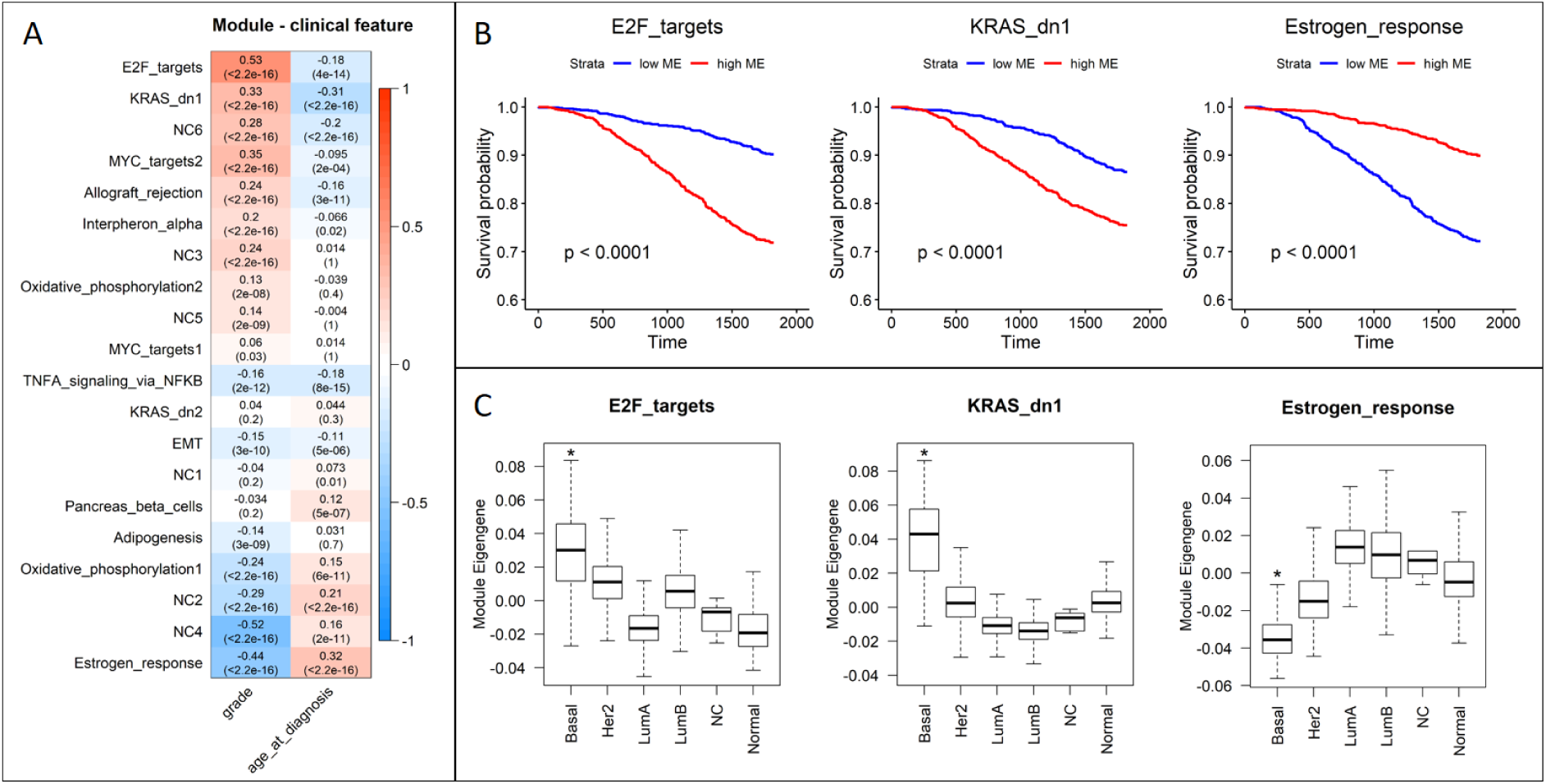
A) Correlation between modules’ expression, represented by module eigengene, and tumour grade and age at diagnosis. Each row corresponds to a different module, columns correspond to clinical features. Red indicates positive and blue negative correlations. For each combination of module-clinical feature, correlation and p-value are shown in the corresponding box. B) E2_targets, KRAS_dn1 and Estrogen_response modules correlate with disease outcome. Patients classified according to the expression levels of the E2_targets and KRAS_dn1 modules display significantly different survival probability, with higher modules expression correlating with poor prognosis. In contrast, Estrogen_response expression significantly correlates with good prognosis. C) E2F_targets, KRAS_dn1 and Estrogen_response modules’ expression across BC molecular subtypes. E2_targets and KRAS_dn1 are significantly up-regulated, while Estrogen_response is significantly down-regulated, in BL tumours. * p-value for the comparison between basal-like versus all other BCs, <2.2*10^−16^.

**Figure 2.**
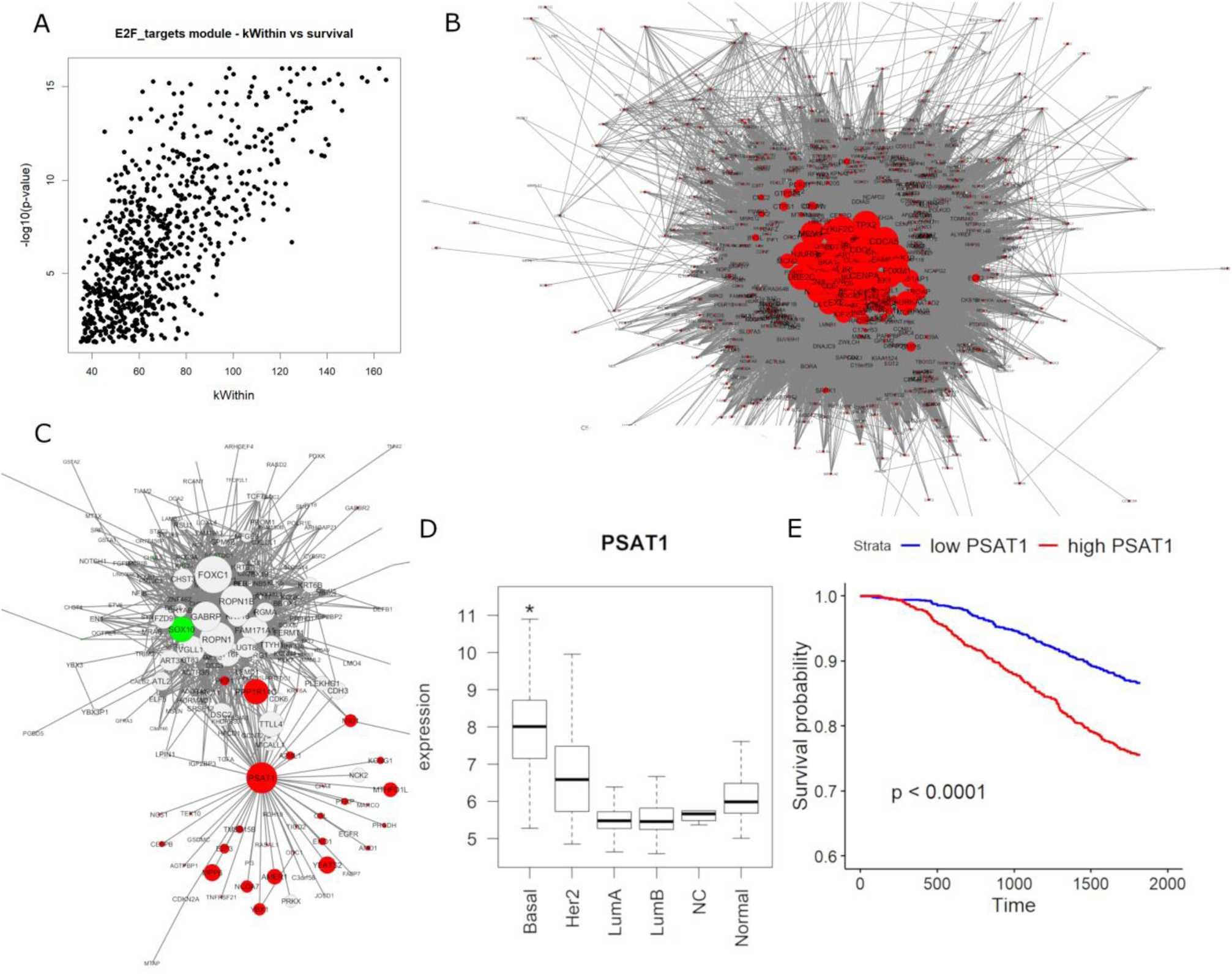
A) Relationship between kWithin of genes in the E2F_targets module and their significance for correlation with poor prognosis. E2F_targets (B) and KRAS_dn1 (C) modules represented with Cytoscape. Each circle is a gene, edges are the strongest topological connections between genes. Red indicates genes correlated with poor prognosis, in green are genes correlated with favourable prognosis. Size of nodes is proportional to kWithin. D) PSAT1 expression across molecular subtypes. * p-value for the comparison between basal-like versus all other subtypes <2.2*10^−16^. E) Survival curves of patients divided according to PSAT1 expression, showing that higher PSAT1 expression significantly correlates with poor prognosis.

### Transcriptional networks are correlated with clinical features

To identify transcriptional networks related with clinical features we calculated for each patient the correlation between modules’ expression and clinical features such as tumour grade and age at diagnosis, as a proxy of age at onset (Fig. 1A). As a measure of module’s expression we used the eigengene, i.e. the ideal gene that best represents the whole module (B. Zhang & Horvath, 2005).

All but three modules showed a significant correlation, either positive or negative, with tumour grade (Fig. 1A), and an opposite trend with respect to age at diagnosis (Table I). In particular, E2F_targets and KRAS_dn1 were the top modules considering the combination of significance between positive correlation with grade and negative correlation with age at diagnosis (Fig. 1A). Thus, tumours with higher expression of either module have features of higher aggressiveness, since they develop earlier and with a higher grade than average (Brandt, Garne, Tengrup, & Manjer, 2015). In contrast, the Estrogen_response module is negatively correlated with grade and positively correlated with age at diagnosis, indicating that its expression is representative of the least aggressive tumours.

**Table 1.** Relationships between module expression and prognosis (5-year disease-specific survival)

| Module | Prognosis | p-value |
| --- | --- | --- |
| E2F_targets | Poor | <2.2*10<sup>-16</sup> |
| MYC_targets2 | Poor | 1.2*10<sup>-15</sup> |
| KRAS_dn1 | Poor | 1*10<sup>-10</sup> |
| NC6 | Poor | 2.2*10<sup>-09</sup> |
| NC3 | Poor | 2*10<sup>-04</sup> |
| NC5 | Poor | 0.0028 |
| Allograft_rejection | Poor | 0.0034 |
| Interferon_alpha | Poor | 0.0058 |
| Oxidative_phosphorylation2 | Poor | 0.014 |
| NC4 | Good | <2.2*10<sup>-16</sup> |
| Estrogen_response | Good | <2.2*10<sup>-16</sup> |
| Oxidative_phosphorylation1 | Good | 1.1*10<sup>-10</sup> |
| NC2 | Good | 1.2*10<sup>-08</sup> |

To further test the relationships with clinical features we calculated the correlation between module eigengenes and patients’ overall survival: both the KRAS_dn1 and E2F_targets modules were significantly associated with poor prognosis, confirming the association between their expression and cancer progression (Fig. 1B). In contrast, high expression of the Estrogen_reponse module was significantly associated with good prognosis. Several other modules were either positively or negatively correlated with survival (Table I, Suppl. Fig. 3,4). Finally, we examined whether modules’ expression was significantly different among breast cancer molecular subtypes (Suppl. Fig. 5), and found that the modules KRAS_dn1 and E2F_targets are the top overexpressed modules in the highly aggressive basal-like tumours (p-values <2.2*10^−16^), while the Estrogen_response module is overexpressed in both luminal subtypes (p-values <2.2*10^−16^) (Fig. 1C, Suppl. Fig. 5). Of note, the E2F_targets module is also overexpressed in the HER2 subtype when compared with all other breast cancer samples (p-value <2.2*10^−16^).

**Figure 3.**
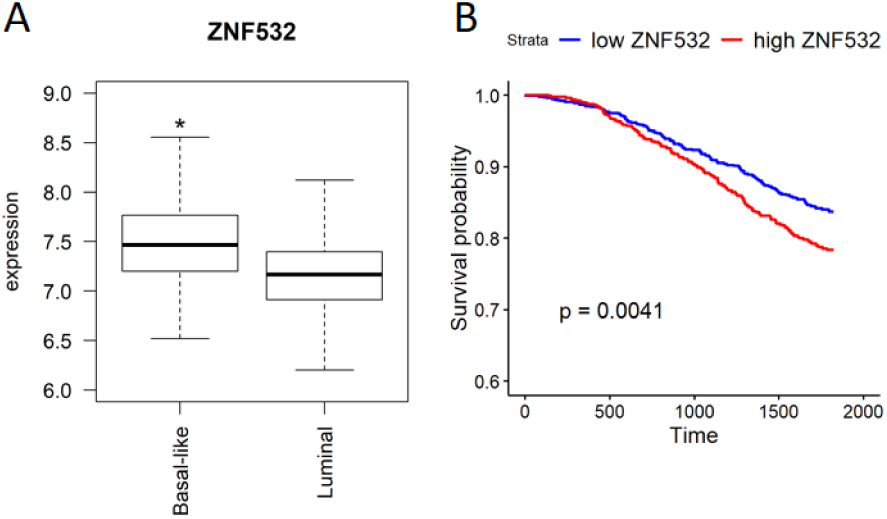
A) ZNF532 expression in basal-like vs luminal (A and B) tumours (* p-value<2.2*10^−16^). B) Survival curves of patients divided based on ZNF532 expression showing that higher ZNF532 expression significantly correlates with poor prognosis.

**Figure 4.**
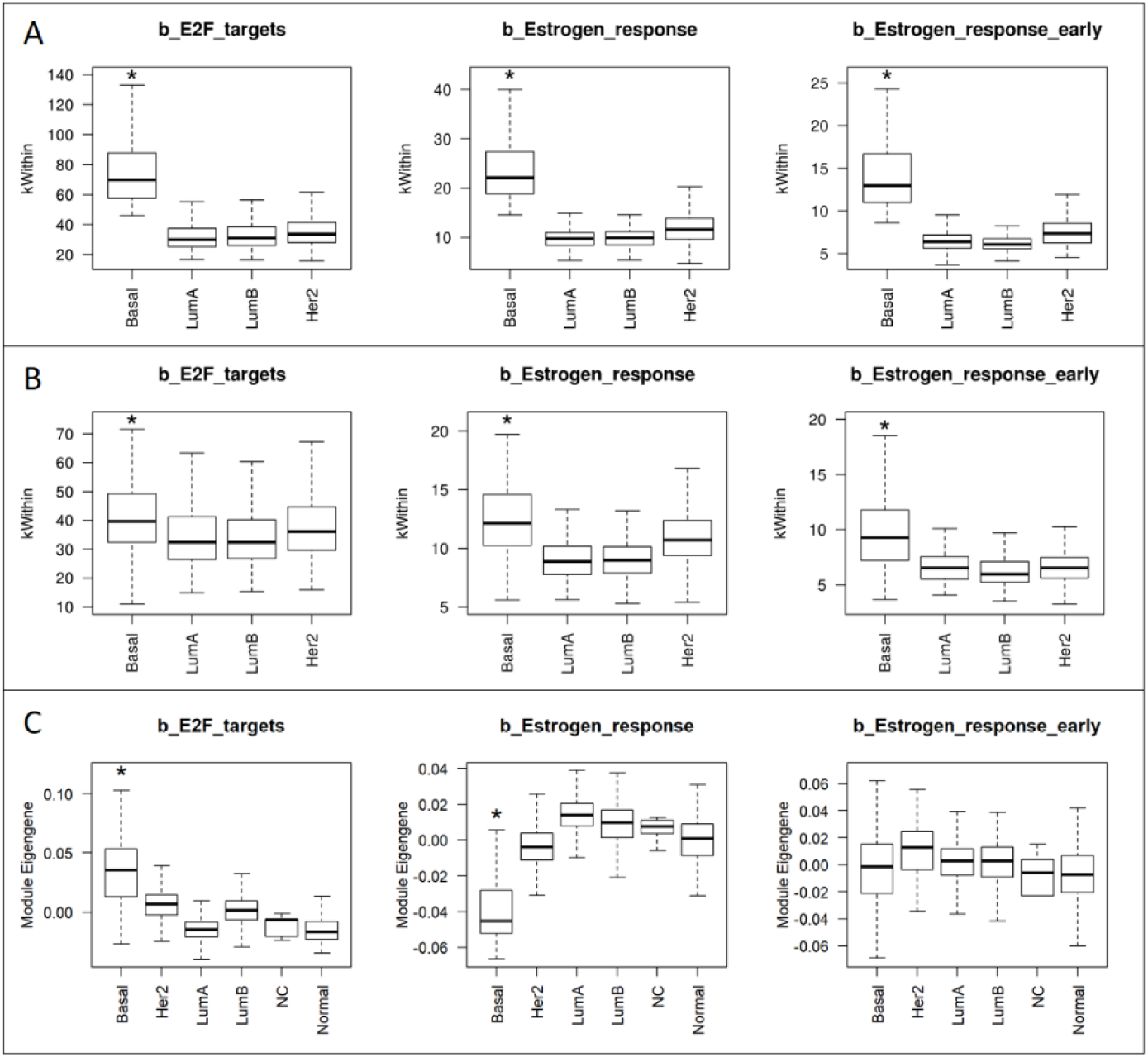
Intramodular connectivity of b_E2F_targets, b_Estrogen_response, b_Estrogen_response_early across BC subtypes as obtained calculating the kWithin in Metabric (A) and in TCGA (B). (C) Expression of the three b_modules represented by Module Eigengene across BC subtypes. * p-value for the comparison of basal-like tumours and all the other subtypes pooled together < 2.2*10^−16^.

**Figure 5.**
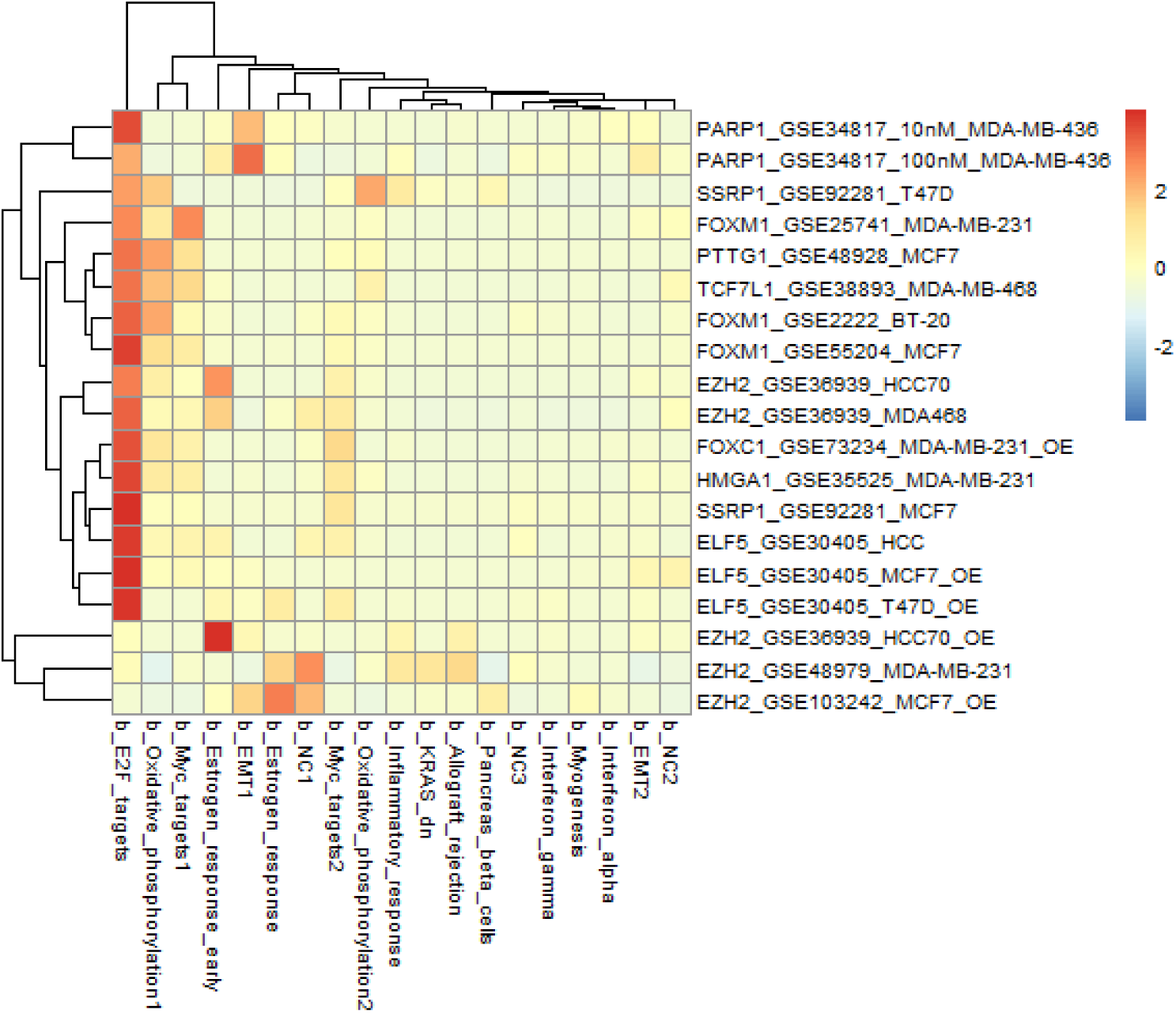
TF hubs in the b_E2F_targets module regulate genes in their same network. Heatmap showing the significance of the enrichment (Fisher test) of differentially expressed genes upon hubs modulation (in the same direction of hubs expression) in modules’ genes. Values are –log10 of p-values, scaled by row. Each row corresponds to a different dataset where a TF has been down-or up-regulated in a BC cell line, as indicated in row names (OE= overexpression). Columns corresponds to different basal modules.

Having thus identified two gene networks of particular interest (E2F_targets and KRAS_dn1), we moved a step forward and defined the most central genes in each module, focusing our interest on transcription factors (TF) as potential candidates for regulating the expression of modules’ genes. As a measure of network centrality we used the intramodular connectivity (kWithin) that we show to be a valuable parameter to identify functionally relevant genes. In fact, the kWithin correlates with the significance of association with poor prognosis for genes in the E2F_targets module (Fig. 2C).

As shown in Fig. 2B, the E2F_targets module’s hubs are cell cycle-related genes, such as CDCA5, TPX2 and CENPA, while the most central TF is FOXM1, scoring as the 14^th^ gene of the module based on the kWithin (Suppl. Table I). Indeed, this TF is well known to be involved in cancer progression and therapy resistance, in particular in BC (reviewed in Saba, Alsayed, Zacny, & Dudek, 2016). The most central genes in the E2F_targets module are overexpressed in basal-like tumours and associated with poor prognosis (Suppl. Fig. 6,7), confirming that they represent well module’s features. The first hub of the KRAS_dn1 module (Fig. 2B) is the FOXC1 TF, well-known to be involved in BC development and metastasis, overexpressed in basal-like breast cancer and associated with poor prognosis (reviewed in Han et al., 2017). The second and third hubs are ROPN1, a cancer-testis antigen involved in sperm motility, and its paralog ROPN1B. Little is known about ROPN1 in breast cancer, but it has been associated with a SOX10 signature shared with salivary adenoid cystic cancer (Ivanov et al., 2013). Interestingly, SOX10 scores as the 8^th^ hub in the same KRAS_dn1 module, and has been reported to be a marker of TNBC (Peevey, Sumpter, Paintal, Laskin, & Sullivan, 2015) and involved both in stem cell activity and EMT in mammary epithelial cells (States et al., 2015). Next is VGLL1, the 6^th^ gene in the kWithin ranking in the KRAS_dn1 module (Suppl. Table I). It is a transcriptional co-activator that binds to several members of the TEAD family of TFs to modulate the Hippo signalling pathway and it is involved in cell cycle regulation. Its association with basal-like BC has only been reported once (Castilla et al., 2014). As observed for the E2F_targets module, also in KRAS_dn1 module the most central genes are overexpressed in the basal-like subtype and often correlated with poor survival (Suppl. Fig. 6,7). Notably, the KRAS_dn1 module is split into two subnetworks connected by a single node, represented by the PSAT1 gene (Phosphoserine aminotransferase 1, involved in serine biosynthesis) (Fig. 2C). Interestingly, one of the two subnetworks is highly enriched in poor prognosis related genes (Suppl. table II), and PSAT1 expression itself correlates with poor prognosis (p-value: 4.5*10^−10^, Fig. 2E), and is significantly enriched in basal-like BC (Fig. 2D). This gene is therefore a good candidate for shaping the KRAS_dn1 module and to coordinate the expression of the module’s genes involved in aggressiveness.

**Table 2.**
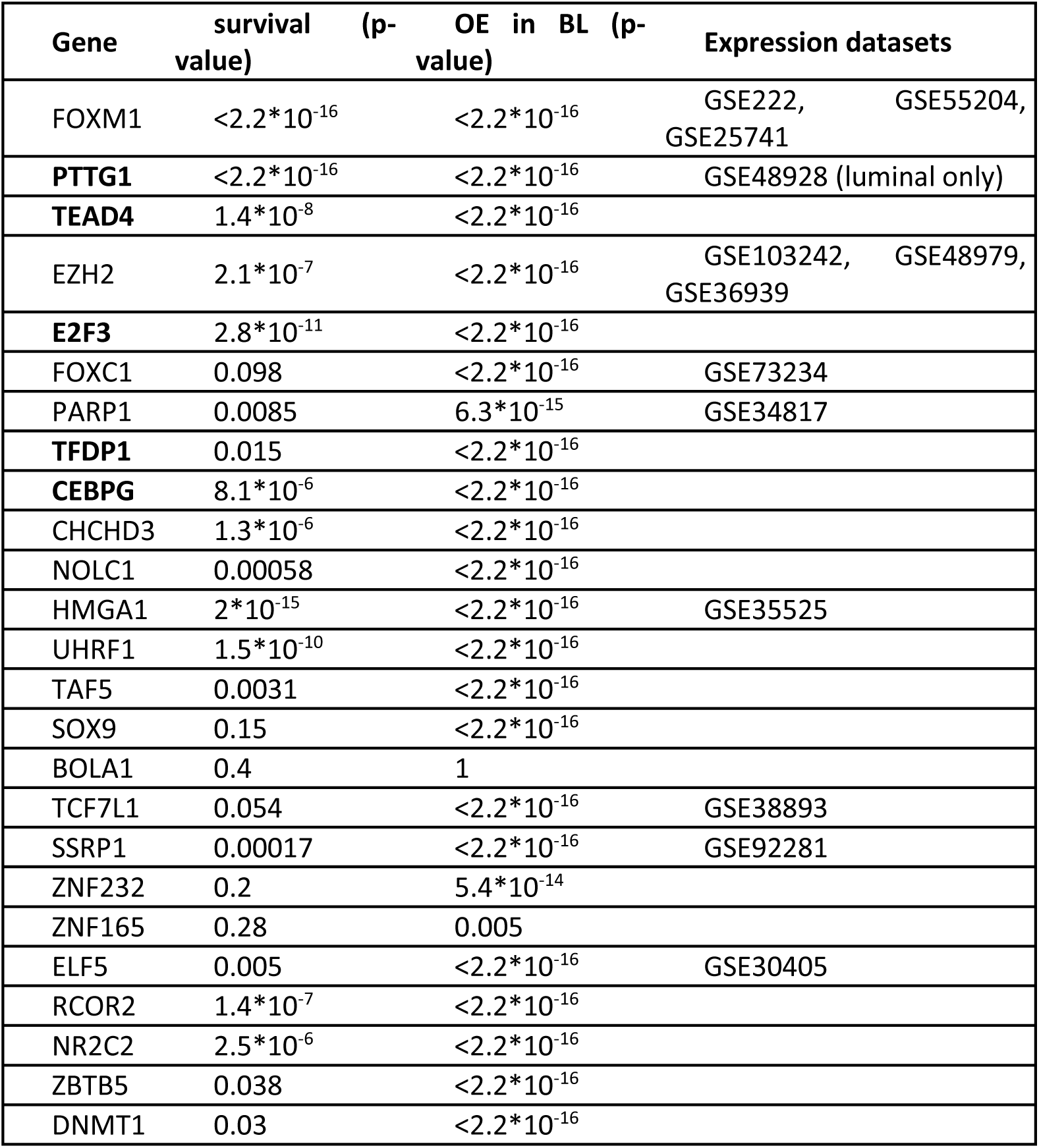
Top centrally located transcription factors in the b_E2F_targets module

**Figure 6.**
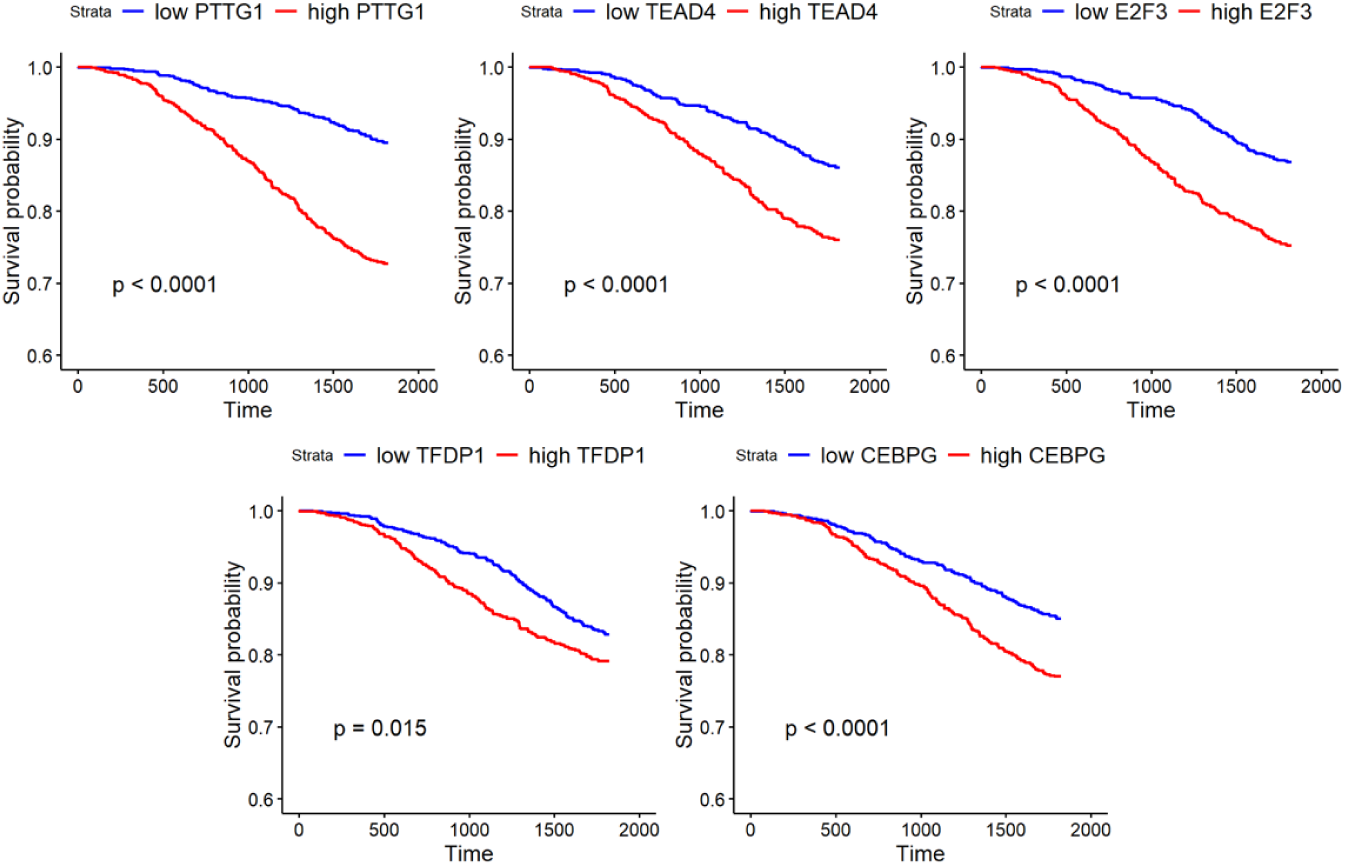
Kaplan-Meier survival curves showing that patients with higher expression of PTTG1, TEAD4, E2F3, TFDP1 or CEBPG display shorter survival.

The Estrogen_response module, overexpressed in luminal tumours, has among its central nodes the TFs GATA3, ESR1 and FOXA1 (respectively 1^st^, 2^nd^ and 7^th^ in the kWIthin ranking, Suppl. Table I), known to be master regulators of ER-positive breast tumours, where they are overexpressed (Suppl. Fig. 6). GATA3 and FOXA1 have both been implicated in shaping the distribution of the ER TF on chromatin and in inhibiting the basal-like phenotype (Bernardo et al., 2012; Hisamatsu, Tokunaga, Yamashita, & Akiyoshi, 2015; Theodorou, Stark, Menon, & Carroll, 2013). Interestingly, amongst the most centrally connected genes are MLPH and P4HTM, whose possible role in BC is poorly characterized.

### ZNF532 is a potential common target of STAT3 and WNT/PCP pathways

As a second approach, we investigated whether the BC networks we constructed could be used to define potential targets of specific pathways. The reasoning behind this idea was that if a gene is transcriptionally activated by a pathway, then it will be co-expressed with genes of the pathway. Therefore pathway genes and potential targets will be part of the same network. Here, we decided to search for potential common targets of the STAT3 and Wnt/PCP pathways, which display several overlapping functions and can interact in a feed-forward loop (Armanious, Gelebart, Mackey, Ma, & Lai, 2010; Gujral et al., 2014; Katoh & Katoh, 2007). The mostly non canonical Wnt5a and Wnt5b ligands are overexpressed in basal-like BC cell lines (Klemm et al., 2011). Moreover, we recently showed that the two pathways converge on the activation of the common downstream target RhoU, promoting cell motility in basal-like human breast tumour cells (Monteleone et al., 2019). Hence, here we aimed at identifying potential additional common targets of the two pathways that could mediate their functions in determining cancer aggressiveness.

As a first step, we collected a set of gene signatures of the two pathways (Alvarez et al., 2005; Azare et al., 2007; Bayerlová et al., 2017; Dauer et al., 2005; Labbe et al., 2007; Sandsmark et al., 2017; Sonnenblick et al., 2015; R W Tell & Horvath, 2014; Willert, Epping, Pollack, Brown, & Nusse, 2002; Ziegler et al., 2005) and we determined which, if any, of the BC modules we defined was enriched in at least one signature of both pathways. Not surprisingly, this analysis pointed to the EMT network, enriched for STAT3 and non-canonical WNT signatures with p-values of 0.001 and 0.02, respectively (Azare et al., 2007; Sandsmark et al., 2017). Via the analysis of a BC ChIP-seq dataset (Mcdaniel et al., 2016), we determined that the EMT module is enriched for genes displaying STAT3 binding at their promoter in the basal-like BC cell lines HCC1143 and MDA-MB-157 (p-values: <0.004 and <3*10^−12^, respectively). Moreover, we analysed an RNA-seq dataset of the basal-like MDA-MB-231 cell line upon STAT3 knock-down, and another one of the luminal cell line MCF7 upon overexpression of the WNT/PCP ROR2 co-receptor (Bayerlová et al., 2017; Mcdaniel et al., 2016). We selected the genes representing the intersection between those down-regulated upon STAT3 knock-down and up-regulated following ROR2 overexpression, hence positively correlated with both STAT3 and WNT/PCP signalling (Suppl. Table III). We then calculated the enrichment of our gene networks for these STAT3/ROR2 regulated genes, finding a significant enrichment for the EMT module (25 genes, p-value=2.15*10^−5^, Suppl. Table IV), followed by the KRAS_dn1 module (p-value=0.001). Therefore, we conclude that the EMT network includes a significant number of genes regulated by both pathways, confirming previous analyses.

Three TFs (ZNF532, RUNX2 and MSX1) are included in these 25 genes (Suppl. Table IV). Interestingly, although the role of ZNF532 in cancer is not known, this factor is overexpressed in basal-like tumours (Fig. 3A), significantly correlating with early tumour onset (Pearson’s correlation=-0.17, p-value=2*10^−14^) and with poor prognosis (p-value: 0.004) (Fig. 3B). This factor represents therefore an interesting candidate as a downstream mediator of the STAT3 and WNT/PCP pathways, potentially involved in regulating all or a subset of the other 24 common target genes identified above.

### Identification of basal-like specific networks

The complexity of reconstructing functionally relevant regulatory networks is partly due to the fact that genes may play multiple functions and that their expression, interactions and significance can be strongly influenced by different conditions, such as, for example, gender or tumour subtype. These potential differences in network structure should be accounted for, and samples belonging to different classes should be separated prior to network reconstruction to avoid misleading inferences, as described in the Simpson Paradox (Simpson, 1951).

To dissect the specificity of gene networks in established molecular subtypes of breast cancer, WGCNA was applied to each set of METABRIC samples classified as basal-like, luminal A, luminal B, HER2-positive. This approach has two advantages: i) having as input a more homogenous set of samples, WGCNA can more reliably identify gene networks based on gene interactions and not on differential gene expression between sets of samples (determined by different molecular subtypes, in this case); ii) it allows identifying differences between network structures across subtypes. 20 gene modules were identified in basal-like tumours, 21 in luminal A, 19 in luminal B, 10 in HER2-positive tumours (not shown). We then focused our interest on basal-like networks, searching for differences in network structure that could account for the higher aggressiveness of this specific subtype. Therefore we selected basal-like specific gene modules based on two criteria: i) at least 40% of module genes fall in the “unconnected” module in other subtypes’ networks (Suppl. Fig. 8); ii) module genes’ intramodular connectivity (kWithin) is higher in basal-like than in other subtypes both in METABRIC, where networks have been defined, and in the independent dataset TCGA (Fig. 4A,B). We did not use gene networks of normal-like tumours due to the low number of samples in TCGA (23) that does not allow a reliable network inference. With this approach we identified three gene modules more tightly co-regulated in basal-like tumours than in the other subtypes. These were named, according to the same criteria used for global modules, b (for basal-like)_E2F_targets, b_Estrogen_response and b_Estrogen_response_early.

Notably, expression of the b_E2F_targets module genes is up-regulated in basal-like tumours while that of the b_Estrogen_response module genes is down-regulated, implying that modules co-regulation and expression levels are independent (Fig. 4C). The b_Estrogen_response module comprises genes specific of ER-positive tumours, such as ESR1 itself, FOXA1 and GATA3, which are known to act in the same network in ER-positive breast tumours (Kong, Li, Loh, Sung, & Liu, 2011). Indeed, the b_Estrogen_response module is enriched for the global Estrogen_response module genes (234/389 genes, Fisher test p-value<2.2*10^−16^), which are in turn up-regulated in luminal molecular subtypes, in agreement with the down-regulation in basal-like tumours (Fig. 4C). A possible interpretation of these results is that luminal specific genes are coherently down-regulated in basal-like tumours, potentially through shared mechanisms, while spreading across different pathways upon activation, thus becoming less coherently regulated.

### Centrally located transcription factors regulate the whole network

We then searched for potential regulatory mechanisms in the identified interesting networks by selecting hubs within each module. Since the undirected networks we generated do not allow to reconstruct upstream and downstream genes in regulatory connections, genes with high kWithin could both be downstream targets of many module genes or upstream regulators. In order to enhance the likelihood of identifying upstream regulators, we specifically searched for transcriptional regulators (TRs), identified as TFs in gene ontology. Strikingly, of the top 25 TRs within the b_E2F_targets module, 75% are correlated with BC poor prognosis and 92% are overexpressed in basal-like tumours (Table II).

We hypothesized that TR hubs of the b_E2F_targets module could be central players in up-regulating the expression of the genes belonging to the network, and might perhaps even be involved in down-regulating the expression of genes belonging to the b_Estrogen_response module in basal-like tumours. To test this hypothesis we searched for published experiments reporting gene expression datasets obtained in BC cell lines upon modulating selected b_E2F_targets TF hubs, in order to assess hubs correlation with global regulation of module’s genes. We found 19 datasets collectively studying 9 different TFs. In 16/19 cases, indeed, genes positively correlated with the selected TFs showed a significant enrichment (p-value<0.001) for genes in the b_E2F_targets module (Fig. 5), validating our hypothesis. The only exception was EZH2, for which contrasting results were obtained in different datasets. Nevertheless, our hypothesis was validated for FOXM1, PTTG1, FOXC1, PARP1, HMGA1, TCF7L1, SSRP1 and ELF5, analysed across several BC cell lines and giving consistent results upon down-and/or up-regulation (Fig. 5). For example, ELF5 knock-down in the basal-like BC cell line HCC1937 leads to the specific down-regulation of b_E2F_targets module genes, while its overexpression in luminal BC cell lines (MCF7, T47D) induces the up-regulation of the same b_E2F_targets module genes. Moreover, we observed that down-regulation of the PTTG1 TF hub induced a set of genes enriched in the b_Estrogen_response module (not shown), suggesting that PTTG1 could be responsible at the same time for b_E2F_targets genes activation and b_Estrogen_response genes repression, thus co-regulating the two networks.

In the light of these results, we expect that additional TFs in a central position within the b_E2F_targets module could orchestrate the expression of the same basal-like specific gene module, therefore likely playing a role in defining the phenotypic features of this subtype. The most central TFs not fully studied in BLBC, all of which correlate with poor prognosis, are PTTG1, TEAD4, E2F3, TFDP1, CEBPG, in bold in Table II (Fig. 6).

## Discussion

Co-expression networks provide a description of gene expression profiles that can be useful to identify sets of functionally related genes. Among the principal aims of network analysis is moreover that of identifying potential regulators of the networks and, consequently, important nodes whose alteration has the potential to disrupt the whole network. Indeed, the number of connections of a gene within a gene expression network was shown to be predictive of its phenotypic relevance in *S. cerevisiae* (Jeong et al., 2001).

Here, we demonstrate that network centrality (kWithin) is a suitable measure to identify relevant genes. In fact, we could show that the kWithin correlates with clinical features such as disease-free survival, and, most importantly, that it provides a mean to select potential upstream regulators of a module with high reliability. As a proof of principle, we combined network measures with the results of independent experiments analyzing gene expression changes induced by altering the expression of selected TF hubs. Strikingly, in 16 out of 19 cases we could show that interference with hub expression induced coherent changes in the expression of their downstream module, demonstrating that indeed our network-based analysis, via the use of modules’ kWithin, provides a valid tool to study biological features of cancer subtypes. In turn this allows to identify potential candidates for networks’ regulation, likely drivers of cancer biological features and useful therapeutic targets. We therefore used the kWithin to prioritize genes and select potential novel candidates for further experimental studies.

We applied our approach to three different issues in breast cancer biology, namely the identification of i) the drivers of basal-like BC aggressiveness; ii) the common targets and potential effectors of the metastasis-related STAT3 and WNT/PCP pathways, and iii) the molecular features distinguishing basal-like BC from the other subtypes.

Our analysis led to the definition of clinically relevant gene modules, such as the E2F_targets and KRAS_dn1 modules, which correlate with BC high grade, early onset, and poor prognosis. Centrally located within these modules are well-known genes involved in BC progression and metastasis such as FOXM1 and FOXC1 (Han et al., 2017; Saba et al., 2016), but also novel uncharacterized genes like ROPN1, involved in sperm motility and whose role in BC has not been studied despite its being associated with the TNBC marker SOX10 (Ivanov et al., 2013; Peevey et al., 2015; States et al., 2015), one of the most central nodes in the same module.

Striking was the observation that the KRAS_dn1 module is divided in two subnetworks, with the PSAT1 gene linking the subnetwork containing the most central hubs (FOXC1, ROPN1) to the subnetwork enriched in poor prognosis-related genes. PSAT1 is the second enzyme in the serine biosynthesis pathway, important for the generation of many cell building blocks (Antonov, Agostini, Morello, & Minieri, 2014; Deberardinis, 2011; Possemato et al., 2011). Several enzymes involved in serine biosynthesis, such as SHMT2 and PHGDH, have been correlated with negative prognosis in BC (Antonov et al., 2014), and PSAT1 itself has been shown to induce proliferation in basal-like BC cell lines (Gao et al., 2017; Marchi et al., 2017). Interestingly, the PSAT1 gene appears to be regulated by FOXC1, a central hub of the same KRAS_dn1 module and associated with BC development (GSE73234, Han et al., 2017). Our data suggest that the correlation of PSAT1 with survival in BC may at least partly be due to its central role in shaping the KRAS_dn1 module.

As a second application, we used our BC networks to define potential targets of specific pathways, and searched for potential common targets of the STAT3 and WNT/PCP pathways. We found evidence of the involvement of the EMT module in STAT3 and WNT/PCP signaling, since this module is significantly enriched in i) both STAT3 and non-canonical WNT signatures; ii) genes whose promoters are bound by STAT3 in several BC cell lines, and iii) genes down-regulated upon STAT3 knock-down in the basal-like cell line MDA-MB-231, and up-regulated upon ROR2 overexpression in the luminal MCF7 cell line. Within the EMT module, the ZNF532 gene, encoding for a TF correlated with poor prognosis in BC (Fig. 3B), appears to be a target of both pathways. We can therefore hypothesize that ZNF532 is a novel common target of the two pathways, and likely a central regulator of the EMT module. This hypothesis can be readily assessed experimentally, and may lead to the identification of a novel target with therapeutic potential, particularly in basal-like BC.

Finally, we dissected the differences in network connectivity across different breast cancer subtypes, comparing the networks obtained with samples of each subtype separately. This analysis is of particular importance when considering that, especially in multicellular organisms, gene networks are context-dependent. Thus, pooling together samples belonging to different groups could result in misleading inferences or hide biologically meaningful signals. We focused on the features distinguishing basal-like networks from networks derived from other subtypes, aiming at identifying molecular mechanisms conferring basal-like BC its aggressiveness and resistance to therapy. Specifically, we defined three modules with stronger intramodular connectivity in the basal-like subtype (Figure 6), two of which show a pattern of differential expression across subtypes, with the b_E2F_targets module being overexpressed in basal-like tumors, and the b_Estrogen_response module being down-regulated in the same subtype. Strikingly, kWithin-based hubs identification and analysis of published gene expression data following their experimental modulation in BC cell lines showed that the most central TFs act indeed as upstream regulators of the module’s genes. This observation represents an empirical validation of the reliability of the kWithin as a selection criterion for hubs that have a high potential of being central regulators of the network structure, and therefore possessing phenotypic relevance. Importantly, among the most central TFs in the b_E2F_targets module are also poor prognosis-related genes for which no experimental studies in BC are available. We thus hypothesize that these represent interesting candidates for phenotypic studies, whose disruption may lead to the dysregulation of the whole basal-like specific module b_E2F_targets, and thus to the inhibition of aggressiveness features of basal-like BC. With these criteria, the genes not yet fully characterized among the top-ranked ones are PTTG1, TEAD4, E2F3, TFDP1, and CEBPG. While PTTG1, TEAD4, E2F3, TFDP1 have been studied to different degrees in BC, C/EBPG was mainly characterized in myeloid tumors (Huggins et al., 2016; Melchor et al., 2009; Vimala, Sundarraj, Sujitha, & Kannan, 2012; Wang et al., 2015; Yoon et al., 2012).

In conclusion, our platform allows to identify groups of genes with a high potential to play a crucial role in basal-like BC and possibly amenable to drug targeting, not only for their specific functions but also for their ability to regulate survival-related gene networks. Finally, our approach could be successfully extended to other BC subtypes, and potentially to all tumor types for which enough expression data are available, allowing to address many additional biological questions.

## METHODS

All analyses were performed using R 3.5.1 and packages obtained from Bioconductor and CRAN (Gentleman et al., 2004; https://cran.r-project.org). Plots were generated with R base graphics, ggplot2 (Wickham, 2017), WGCNA (Langfelder & Horvath, 2008) and pheatmap.

### Network construction

Gene co-expression networks were constructed with WGCNA function blockwiseModules (Langfelder & Horvath, 2008), using as parameters: power= 6, corType=“pearson”, networkType = “signed”, minModuleSize = 30, reassignThreshold = 0, mergeCutHeight = 0.25. METABRIC gene expression data and metadata were obtained from www.synapse.org (syn2133318, syn2133322, syn2133500). Probe names were converted in Gene Symbols and, for gene symbols corresponding to multiple probes, the most expressed probe across all samples was considered.

Preservation was obtained with the function modulePreservation in WGCNA using, independently, breast primary tumors data and BC cell lines data (http://cancergenome.nih.gov/, Daemen et al., 2013). STRING was analysed with the package STRINGdb.

### Correlation with clinical features

Module eigengene calculated on the whole dataset (Langfelder & Horvath, 2008) was used as a measure modules’ expression in each sample. Pearson’s correlation between module eigengene and either grade or age at diagnosis was calculated and p-values were obtained with the function corPvalueFisher (Fisher’s asymptotic p-value). P-values were adjusted using the Holm method in the function p.adjust.

Differences in gene or modules’ expression across subtypes were assessed via a two-sided t-test on z-scores (for individual genes) or on module eigengenes (for modules). Expression in one subtype was compared with the expression in all other subtypes pooled together.

### Correlation with survival

The correlation between the expression level of a gene or a module was calculated dividing the patients in two groups based on the gene or module’s expression median. The function survdiff of the survival package (Mohamed, Abdelaal, Hossam, & Ahmed, 2015), implementing a log-rank statistic, was used and Kaplan-Meier curves were plotted with the ggsurvplot function in the survminer package.

### STAT3 and WNT signatures

A set of STAT3 and WNT signatures was collected (Alvarez et al., 2005; Azare et al., 2007; Bayerlová, Klemm, Kramer, & Pukrop, 2015; Dauer et al., 2005; Labbe et al., 2007; Sandsmark et al., 2017; Sonnenblick et al., 2015; Robert W Tell & Horvath, 2014; Willert et al., 2002; Ziegler et al., 2005) and individual enrichments of modules for each signature were calculated with a Fisher’s exact test using as using as background the list of genes not falling in the “unconnected” compartment. P-values were adjusted with the p.adjust function and the Holm method considering all the tests performed.

Differentially expressed genes upon STAT3 down-regulation in MDA-MB-231 were defined from Mcdaniel et al., 2016 using DESeq2 (Love, Huber, & Anders, 2014) and a p-value cutoff of 0.05. Differentially expressed genes upon ROR2 overexpression in MCF7 were obtained from Bayerlová et al., 2017 with p-value<0.05.The enrichment of modules for genes in the intersection of these lists of differentially expressed genes was calculated with a Fisher’s exact test.

### ChIP-seq data analysis

ChIP-seq data (Mcdaniel et al., 2016) were mapped to the H. sapiens hg19 genome build with bowtie 1.2.1 (parameters: -p 3 –best –strata –m1 –S, Langmead, Trapnell, Pop, & Salzberg, 2009). Peaks were defined with MACS 1.4.2 (Y. Zhang et al., 2008) and the closest genes to peaks were annotated with the function closestBed (BEDTools suite, Quinlan & Hall, 2010). Peaks with score ≥ 200 were retained, and those in the range between −1000bp and 0bp from the transcription start site were considered indicative of binding at the promoter. The enrichment for genes in each module was calculated with Fisher’s exact test.

### Subtype-specific modules

Subtype-specific networks were constructed on basal-like, luminal A, luminal B and HER2-positive tumour samples separately using WGCNA with the same parameters used for the global network. Comparisons of intramodular connectivity (kWithin) were performed as follows: kWithin were calculated on all combinations of subtype-specific networks and assignments of genes to modules to determine the connectivity of genes of a module, as defined in a subtype, in the other subtypes; distributions of kWithin of genes of a module were compared across subtypes using a t-test.

### Analysis of TF datasets

Pre-processed gene expression data were downloaded from Gene Expression Omnibus (GSE2222, GSE55204, GSE25741, GSE48928, GSE103242, GSE48979, GSE36939, GSE73234, GSE34817, GSE35525, GSE38893, GSE92281, GSE30405). Lists of differentially expressed genes were obtained with a t-test (function t.test) and a p-value cutoff of 0.05. Enrichment of differentially expressed genes for modules’ genes were calculated with a Fisher’s exact test using as background the list of genes not falling in the “unconnected” compartment in basal-like tumors and which expression could be retrieved based on the specific array used in the dataset.

### Networks representations

The most strongly connected nodes (Topological Overlap ≥ 0.02) were retained for individual modules’ visualization in Cytoscape 3.7.0 (Shannon et al., 2003). The whole network was filtered for connections with Topological Overlap ≥ 0.05 and represented with Gephi 0.9.2 (Bastian & Heymann, 2009).

## Supporting information

Supplementary Table III

## ACKNOWLEDGMENTS

This work was supported by the Italian Cancer Research Association (AIRC IG16930 to V.P.) and the Truus and Gerrit van Riemsdijk Foundation, Liechtenstein, donation to V.P.

## References

Albert, R., Jeong, H., & Barabási, A.-L. (2000). Error and attack tolerance of complex networks. Nature, 406, 378. https://doi.org/10.1038/35019019

Alvarez, J. V, Febbo, P. G., Ramaswamy, S., Loda, M., Richardson, A., & Frank, D. A. (2005). Identification of a Genetic Signature of Activated Signal Transducer and Activator of Transcription 3 in Human Tumors. Cancer Research, 65(12), 5054–5063.

Antonov, A., Agostini, M., Morello, M., & Minieri, M. (2014). Bioinformatics analysis of the serine and glycine pathway in cancer cells. Oncotarget, 5(22).

Armanious, H., Gelebart, P., Mackey, J., Ma, Y., & Lai, R. (2010). STAT3 upregulates the protein expression and transcriptional activity of β-catenin in breast cancer. International Journal of Clinical Experimental Pathology, 3(7), 654–664.

Azare, J., Leslie, K., Al-Ahmadie, H., Gerald, W., Weinreb, P. H., Violette, S. M., & Bromberg, J. (2007). Constitutively activated Stat3 induces tumorigenesis and enhances cell motility of prostate epithelial cells through integrin beta 6. Molecular and Cellular Biology, 27(12), 4444–4453. https://doi.org/10.1128/MCB.02404-06

Barabási, A., & Oltvai, Z. N. (2004). Network biology: understanding the cell’s functional organization. Nature Reviews Genetics, 5. https://doi.org/10.1038/nrg1272

Bastian, M., & Heymann, S. (2009). Gephi : An Open Source Software for Exploring and Manipulating Networks. AAAI.

Bayerlová, M., Klemm, F., Kramer, F., & Pukrop, T. (2015). Newly Constructed Network Models of Different WNT Signaling Cascades Applied to Breast Cancer Expression Data. PLoS ONE, 10(12), 1–19. https://doi.org/10.1371/journal.pone.0144014

Bayerlová, M., Menck, K., Klemm, F., Wolff, A., Pukrop, T., Gutenberg-universität, J., … Beißbarth, T. (2017). Ror2 Signaling and Its Relevance in Breast Cancer Progression. Frontiers in Oncology, 7, 1–16. https://doi.org/10.3389/fonc.2017.00135

Bernardo, G. M., Bebek, G., Ginther, C. L., Sizemore, S. T., Lozada, K. L., Miedler, J. D., … Abdul-karim, F. W. (2012). FOXA1 represses the molecular phenotype of basal breast cancer cells. Oncogene, 32(5), 554– 563. https://doi.org/10.1038/onc.2012.62

Brandt, J., Garne, J. P., Tengrup, I., & Manjer, J. (2015). Age at diagnosis in relation to survival following breast cancer?: a cohort study. World Journal of Surgical Oncology, 13(33), 1–11. https://doi.org/10.1186/s12957-014-0429-x

Chasman, D., Siahpirani, A. F., & Roy, S. (2016). ScienceDirect Network-based approaches for analysis of complex biological systems. Current Opinion in Biotechnology, 39, 157–166. https://doi.org/10.1016/j.copbio.2016.04.007

Clarke, C., Madden, S. F., Doolan, P., Aherne, S. T., Joyce, H., Driscoll, L. O., … Clynes, M. (2013). Correlating transcriptional networks to breast cancer survival?: a large-scale coexpression analysis. Carcinogenesis, 34(10), 2300–2308. https://doi.org/10.1093/carcin/bgt208

Curtis, C., Shah, S. P., Chin, S.-F., Turashvili, G., Rueda, O. M., Dunning, M. J., … Aparicio, S. (2012). The genomic and transcriptomic architecture of 2,000 breast tumours reveals novel subgroups. Nature, 486, 346. http://dx.doi.org/10.1038/nature10983

Dauer, D. J., Ferraro, B., Song, L., Yu, B., Mora, L., Buettner, R., … Haura, E. B. (2005). Stat3 regulates genes common to both wound healing and cancer. Oncogene, 24, 3397–3408. https://doi.org/10.1038/sj.onc.1208469

Deberardinis, R. J. (2011). Previews Serine Metabolism : Some Tumors Take the Road Less Traveled. Cell Metabolism, 14(3), 285–286. https://doi.org/10.1016/j.cmet.2011.08.004

Emilsson, V., Thorleifsson, G., Zhang, B., Leonardson, A. S., Zink, F., Zhu, J., … Stefansson, K. (2008). Genetics of gene expression and its effect on disease. Nature, 452, 423. https://doi.org/10.1038/nature06758

Feldman, I., Rzhetsky, A., & Vitkup, D. (2008). Network properties of genes harboring inherited disease mutations. Proceedings of the National Academy of Sciences, 105(11).

Furlong, L. I. (2013). Human diseases through the lens of network biology. Trends in Genetics, 29(3), 150–159. https://doi.org/10.1016/j.tig.2012.11.004

Gao, S., Ge, A., Xu, S., You, Z., Ning, S., Zhao, Y., & Pang, D. (2017). PSAT1 is regulated by ATF4 and enhances cell proliferation via the GSK3 ß / ß-catenin / cyclin D1 signaling pathway in ER-negative breast cancer. Journal of Experimental & Clinical Cancer Research, 36(179), 1–13. https://doi.org/10.1186/s13046-017-0648-4

Gentleman, R. C., Carey, V. J., Bates, D. M., Bolstad, B., Dettling, M., Dudoit, S., … Zhang, J. (2004). Bioconductor?: open software development for computational biology and bioinformatics. Genome Biology, 5(10).

Goh, K., Cusick, M. E., Valle, D., Childs, B., & Vidal, M. (2007). The human disease network. Proceedings of the National Academy of Sciences, 104(21), 8685–8690.

Gujral, T. S., Chan, M., Peshkin, L., Sorger, P. K., Kirschner, M. W., & Macbeath, G. (2014). Article A Noncanonical Frizzled2 Pathway Regulates Epithelial-Mesenchymal Transition and Metastasis. Cell, 159(4), 844–856. https://doi.org/10.1016/j.cell.2014.10.032

Han, B., Bhowmick, N., Qu, Y., Chung, S., Giuliano, A. E., & Cui, X. (2017). FOXC1: an emerging marker and therapeutic target for cancer. Oncogene, 36, 3957–3963. https://doi.org/10.1038/onc.2017.48

Hartwell, L. H., Hopfield, J. J., Leibler, S., & Murray, A. W. (1999). From molecular to modular cell biology. Nature, 402, C47. https://doi.org/10.1038/35011540

Herschkowitz, J. I., Komurov, K., Zhou, A. Y., Gupta, S., Yang, J., Hartwell, K., … Yang, J. (2010). Core epithelial-to-mesenchymal transition interactome gene-expression signature is associated with claudin-low and metaplastic breast cancer subtypes. Proceedings of the National Academy of Sciences, 107(44), 19132. https://doi.org/10.1073/pnas.1015095107

Hisamatsu, Y., Tokunaga, E., Yamashita, N., & Akiyoshi, S. (2015). Impact of GATA-3 and FOXA1 expression in patients with hormone receptor-positive/HER2-negative breast cancer. Breast Cancer, 22(5), 520–528. https://doi.org/10.1007/s12282-013-0515-x

Huggins, C. J., Mayekar, M. K., Martin, N., Saylor, K. L., Gonit, M., Jailwala, P., … Johnson, F. (2016). C/EBPg Is a Critical Regulator of Cellular Stress Response Networks through Heterodimerization with ATF4. Molecular and Cellular Biology, 36(5), 693–713. https://doi.org/10.1128/MCB.00911-15.Address

Isik, Z., Baldow, C., Cannistraci, C. V., & Schroeder, M. (2015). Drug target prioritization by perturbed gene expression and network information. Scientific Reports, 5, 1–13. https://doi.org/10.1038/srep17417

Ivanov, S. V, Panaccione, A., Nonaka, D., Prasad, M. L., Boyd, K. L., Brown, B., … Yarbrough, W. G. (2013). Diagnostic SOX10 gene signatures in salivary adenoid cystic and breast basal-like carcinomas. British Journal of Cancer, 109(2), 444–51. https://doi.org/10.1038/bjc.2013.326

Jensen, L. J., Kuhn, M., Stark, M., Chaffron, S., Creevey, C., Muller, J., … von Mering, C. (2009). STRING 8--a global view on proteins and their functional interactions in 630 organisms. Nucleic Acids Research, 37(Database issue), D412–D416. https://doi.org/10.1093/nar/gkn760

Jeong, H., Mason, S. P., Barabási, A.-L., & Oltvai, Z. N. (2001). Lethality and centrality in protein networks. Nature, 411, 41. https://doi.org/10.1038/35075138

Katoh, M., & Katoh, M. (2007). STAT3-induced WNT5A signaling loop in embryonic stem cells, adult normal tissues, chronic persistent inflammation, rheumatoid arthritis and cancer. International Journal of Molecular Medicine, 19(2), 273–278.

Klein, K. O., Oualkacha, K., Lafond, M., & Bhatnagar, S. (2016). Gene Coexpression Analyses Differentiate Networks Associated with Diverse Cancers Harboring TP53 Missense or Null Mutations. Frontiers in Genetics, 7, 1–14. https://doi.org/10.3389/fgene.2016.00137

Klemm, F., Bleckmann, A., Siam, L., Chuang, H. N., Rietktter, E., Behme, D., … Pukrop, T. (2011). ßcatenin-independent WNT signaling in basal-like breast cancer and brain metastasis. Carcinogenesis, 32(3), 434–442. https://doi.org/10.1093/carcin/bgq269

Labbe, E., Lock, L., Letamendia, A., Gorska, A. E., Gryfe, R., Gallinger, S., … Attisano, L. (2007). Transcriptional Cooperation between the Transforming Growth Factor-B and Wnt Pathways in Mammary and Intestinal Tumorigenesis. Cancer Research, 67(1), 75–85. https://doi.org/10.1158/0008-5472.CAN-06-2559

Langfelder, P., & Horvath, S. (2008). WGCNA: an R package for weighted correlation network analysis. BMC Bioinformatics, 9, 559. https://doi.org/10.1186/1471-2105-9-559

Langmead, B., Trapnell, C., Pop, M., & Salzberg, S. L. (2009). Ultrafast and memory-efficient alignment of short DNA sequences to the human genome. Genome Biology, 10(3). https://doi.org/10.1186/gb-2009-10-3-r25

Liu, L., Chen, X., Hu, C., Zhang, D., Shao, Z., Jin, Q., … Xie, H. (2018). Synthetic Lethality-based Identification of Targets for Anticancer Drugs in the Human Signaling Network. Scientific Reports, 8(8440), 1–10. https://doi.org/10.1038/s41598-018-26783-w

Liu, X., Bulgakov, O. V, Darrow, K. N., Pawlyk, B., Adamian, M., Liberman, M. C., & Li, T. (2007). Usherin is required for maintenance of retinal photoreceptors and normal development of cochlear hair cells. Proceedings of the National Academy of Sciences, 104(11), 4413–4418. https://doi.org/10.1073/pnas.0610950104

Love, M. I., Huber, W., & Anders, S. (2014). Moderated estimation of fold change and dispersion for RNA-seq data with DESeq2. Genome Biology, 15(12), 550. https://doi.org/10.1186/s13059-014-0550-8

Malod-dognin, N., Petschnigg, J., Windels, S. F. L., Povh, J., Hemmingway, H., & Ketteler, R. (2019). Towards a data-integrated cell. Nature Communications, 10(805), 1–13. https://doi.org/10.1038/s41467-019-08797-8

Marchi, T. De, Timmermans, M. A., Sieuwerts, A. M., Smid, M., Look, M. P., Greben, N., … Martens, J. W. (2017). Phosphoserine aminotransferase 1 is associated to poor outcome on tamoxifen therapy in recurrent breast cancer. Scientific Reports, (7), 1–11. https://doi.org/10.1038/s41598-017-02296-w

Mcdaniel, J. M., Varley, K. E., Gertz, J., Savic, D. S., Brian, S., Bailey, S. K., … Myers, R. M. (2016). Genomic regulation of invasion by STAT3 in triple negative breast cancer. Oncotarget, 8(5).

Melchor, L., Saucedo-cuevas, L. P., Muñoz-repeto, I., Rodríguez-pinilla, M., Honrado, E., Campoverde, A., … Benítez, J. (2009). Research article Comprehensive characterization of the DNA amplification at 13q34 in human breast cancer reveals TFDP1 and CUL4A as likely candidate target genes. Breast Cancer Research, 11(6), 1–14. https://doi.org/10.1186/bcr2456

Mohamed, M., Abdelaal, A., Hossam, S., & Ahmed, E. (2015). Modeling Survival Data by Using Cox Regression Model. American Journal of Theoretical and Applied Statistics, 4(6), 504–512. https://doi.org/10.11648/j.ajtas.20150406.21

Monteleone, E., Orecchia, V., Corrieri, P., Schiavone, D., Avalle, L., Moiso, E., … Poli, V. (2019). SP1 and STAT3 Functionally Synergize to Induce the RhoU Small GTPase and a Subclass of Non-canonical WNT Responsive Genes Correlating with Poor Prognosis in Breast Cancer. Cancers, 11(1), 101–116. https://doi.org/10.3390/cancers11010101

Peevey, J., Sumpter, I., Paintal, A., Laskin, W., & Sullivan, M. (2015). SOX10 Is a Useful Marker for Triple Negative Breast Cancer. American Journal of Clinical Pathology, 144(Suppl_2), A299–A299. http://dx.doi.org/10.1093/ajcp/144.suppl2.299

Perou, C. M., Sørlie, T., Eisen, M. B., van de Rijn, M., Jeffrey, S. S., Rees, C. A., … Botstein, D. (2000). Molecular portraits of human breast tumours. Nature, 406, 747. https://doi.org/10.1038/35021093

Possemato, R., Marks, K. M., Shaul, Y. D., Pacold, M. E., Kim, D., Birsoy, K., … Sabatini, D. M. (2011). pathway is essential in breast cancer. Nature, 476(7360), 346–350. https://doi.org/10.1038/nature10350

Quinlan, A. R., & Hall, I. M. (2010). BEDTools?: a flexible suite of utilities for comparing genomic features. Bioinformatics, 26(6), 841–842. https://doi.org/10.1093/bioinformatics/btq033

Saba, R., Alsayed, A., Zacny, J. P., & Dudek, A. Z. (2016). The Role of Forkhead Box Protein M1 in Breast Cancer Progression and Resistance to Therapy. International Journal of Breast Cancer, 2016. http://dx.doi.org/10.1155/2016/9768183

Saelens, W., Cannoodt, R., & Saeys, Y. (2018). A comprehensive evaluation of module detection methods for gene expression data. Nature Communications, 9(1090). https://doi.org/10.1038/s41467-018-03424-4

Sandsmark, E., Hansen, A. F., Selnæs, K. M., Bertilsson, H., Bofin, A. M., Wright, A. J., … Rye, M. B. (2017). A novel non-canonical Wnt signature for prostate cancer aggressiveness. Oncotarget, 8(6), 9572–9586.

Shannon, P., Markiel, A., Ozier, O., Baliga, N. S., Wang, J. T., Ramage, D., … Ideker, T. (2003). Cytoscape?: A Software Environment for Integrated Models of Biomolecular Interaction Networks. Genome Research, 13(11), 2498–2504. https://doi.org/10.1101/gr.1239303.metabolite

Shi, H., Zhang, L. E. I., Qu, Y., Hou, L., Wang, L., & Zheng, M. I. N. (2017). Prognostic genes of breast cancer revealed by gene co-expression network analysis. Oncology Letters, 14, 4535–4542. https://doi.org/10.3892/ol.2017.6779

Simpson, E. H. (1951). The Interpretation of Interaction in Contingency Tables. Journal of the Royal Statistical Society. Series B (Methodological), 13(2), 238–241. http://www.jstor.org/stable/2984065

Sonnenblick, A., Brohée, S., Fumagalli, D., Vincent, D., Venet, D., Ignatiadis, M., … Sotiriou, C. (2015). Constitutive phosphorylated STAT3-associated gene signature is predictive for trastuzumab resistance in primary HER2-positive breast cancer. BMC Medicine, 13(1), 177. https://doi.org/10.1186/s12916-015-0416-2

States, C., Cells, E., Dravis, C., Spike, B. T., Harrell, J. C., Smith, E. M. S.-, … Wahl, G. M. (2015). Sox10 Regulates Stem/Progenitor and Mesenchymal Article Sox10 Regulates Stem/Progenitor and Mesenchymal Cell States in Mammary Epithelial Cells. Cell Reports, 12(12), 2035–2048. https://doi.org/10.1016/j.celrep.2015.08.040

Tell, R. W., & Horvath, C. M. (2014). Bioinformatic analysis reveals a pattern of STAT3-associated gene expression specific to basal-like breast cancers in human tumors. Proc Natl Acad Sci U S A, 111(35), 12787– 12792. https://doi.org/10.1073/pnas.1404881111

Theodorou, V., Stark, R., Menon, S., & Carroll, J. S. (2013). GATA3 acts upstream of FOXA1 in mediating ESR1 binding by shaping enhancer accessibility. Genome Research, 23(1), 12–22. https://doi.org/10.1101/gr.139469.112.12

Vimala, K., Sundarraj, S., Sujitha, M. V, & Kannan, S. (2012). Curtailing Overexpression of E2F3 in Breast Cancer Using siRNA(E2F3)-Based Gene Silencing. Archives of Medical Research, 43(6), 415–422. https://doi.org/10.1016/j.arcmed.2012.08.009

Wang, C., Nie, Z., Zhou, Z., Zhang, H., Liu, R., Qin, J., … Chen, C. (2015). The interplay between TEAD4 and KLF5 promotes breast cancer partially through inhibiting the transcription of p27 Kip1. Oncotarget, 6(19).

Wickham, H. (2017). ggplot2 – Elegant Graphics for Data Analysis. Journal of Statistical Software April, 77, 3–5. https://doi.org/10.18637/jss.v077.b02

Willert, J., Epping, M., Pollack, J. R., Brown, P. O., & Nusse, R. (2002). A transcriptional response to Wnt protein in human embryonic carcinoma cells. BMC Developmental Biology, 2(8), 1–7.

Wolf, D. M., Lenburg, M. E., Yau, C., Boudreau, A., & Van Veer, L. J.. (2014). Gene Co-Expression Modules as Clinically Relevant Hallmarks of Breast Cancer Diversity. PLoS ONE, 9(2). https://doi.org/10.1371/journal.pone.0088309

Yamanishi, Y., Araki, M., Gutteridge, A., Honda, W., & Kanehisa, M. (2008). Prediction of drug – target interaction networks from the integration of chemical and genomic spaces. Bioinformatics, 24, 232–240. https://doi.org/10.1093/bioinformatics/btn162

Yang, Y., Han, L., Yuan, Y., Li, J., Hei, N., & Liang, H. (2014). Gene co-expression network analysis reveals common system-level properties of prognostic genes across cancer types. Nature Communications, 5(3231), 1–9. https://doi.org/10.1038/ncomms4231

Yoon, C., Kim, M., Lee, H., Kim, R., Lim, E., Yoo, K., … Lee, S. (2012). PTTG1 Oncogene Promotes Tumor Malignancy via Epithelial to Mesenchymal Transition and Expansion of Cancer Stem. Journal of Biological Chemistry, 287(23), 19516–19527. https://doi.org/10.1074/jbc.M111.337428

Yu, W., Zhao, S., Wang, Y., Zhao, B. N., & Zhao, W. (2017). Identification of cancer prognosis-associated functional modules using differential co-expression networks. Oncotarget, 8(68), 112928–112941.

Zhang, B., & Horvath, S. (2005). A General Framework for Weighted Gene Co-Expression Network Analysis. Statistical Applications in Genetics and Molecular Biology, 4(1). https://doi.org/10.2202/1544-6115.1128

Zhang, Y., Liu, T., Meyer, C. A., Eeckhoute, J., Johnson, D. S., Bernstein, B. E., … Liu, X. S. (2008). Model-based Analysis of ChIP-Seq (MACS). Genome Biology, (9). https://doi.org/10.1186/gb-2008-9-9-r137

Zhao, W., Langfelder, P., Fuller, T., Dong, J., Li, A., Hovarth, S., … Hovarth, S. (2010). Weighted Gene Coexpression Network Analysis: State of the Art. Journal of Biopharmaceutical Statistics, 20(2). https://doi.org/10.1080/10543400903572753

Zhu, L., Ding, Y., Chen, C., Wang, L., Huo, Z., Kim, S., … Tseng, G. C. (2017). MetaDCN?: meta-analysis framework for differential co-expression network detection with an application in breast cancer. Bioin, 33(8), 1121–1129. https://doi.org/10.1093/bioinformatics/btw788

Ziegler, S., Ro, S., Tickenbrock, L., Mo, T., Vetter, I. R., & Mu, O. (2005). Novel target genes of the Wnt pathway and statistical insights into Wnt target promoter regulation. FEBS Journal, 272, 1600–1615. https://doi.org/10.1111/j.1742-4658.2005.04581.x

