## Supplementary Figures for "Network analysis allows to unravel breast cancer molecular features and to identify novel targets"

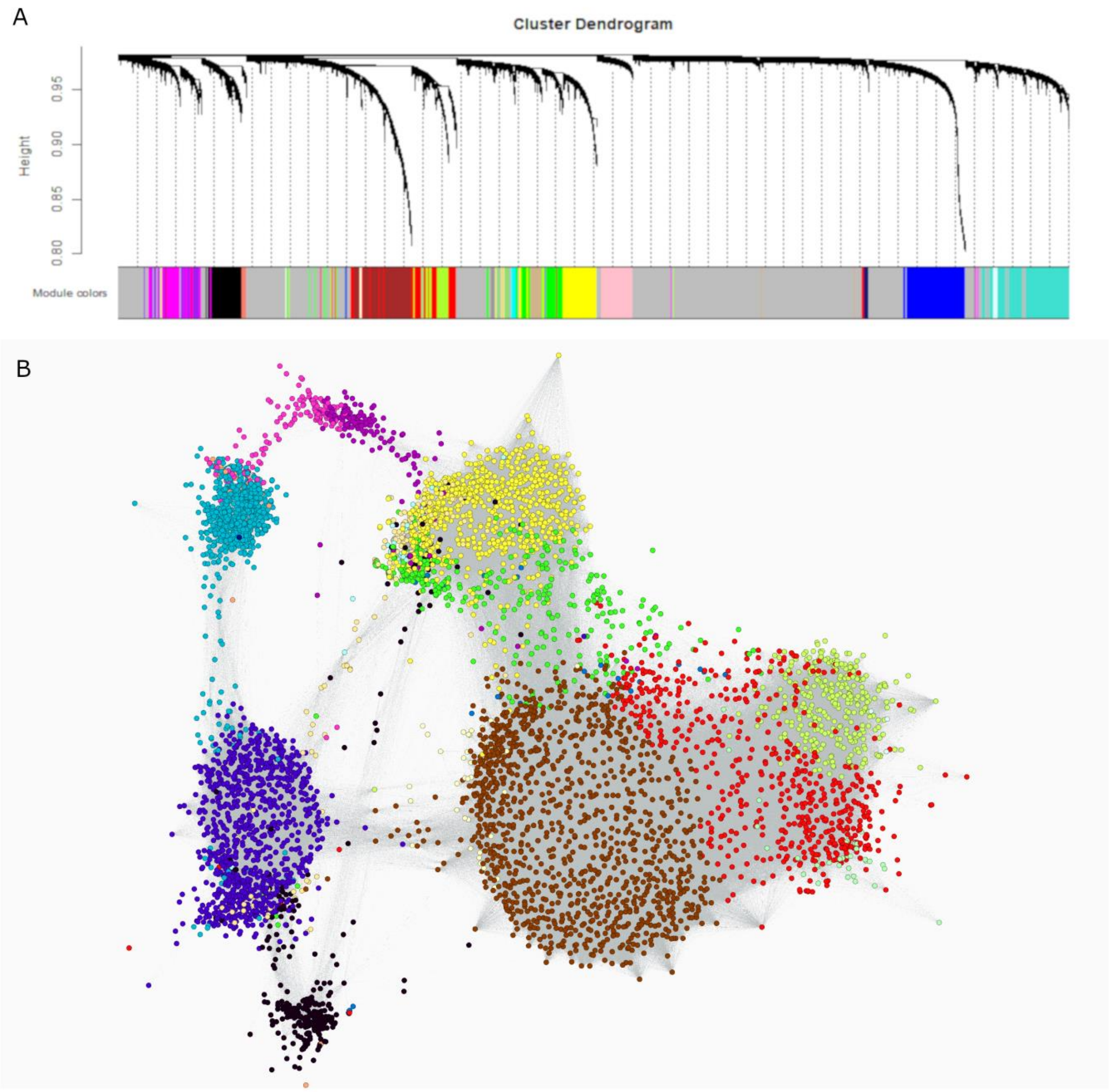


**Supplementary Figure 1.** A) Clustering of genes performed with WGCNA, each colour corresponds to a module. B) Gene co-expression network represented with Gephi: nodes are genes and edges are represented only for the strongest connections (Topological Overlap ≥ 0.05) in the network. Different colours correspond to different modules.

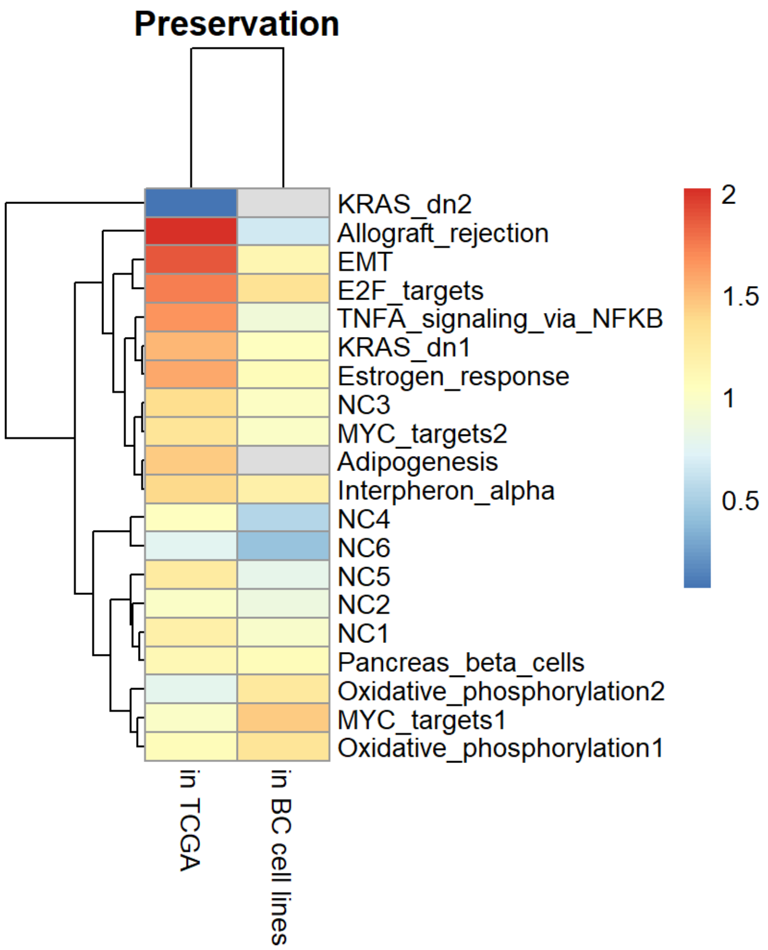


**Supplementary Figure 2.** Preservation Zscores (log10) of METABRIC modules in TCGA and in BC cell lines (http://cancergenome.nih.gov/, Daemen et al., 2013)

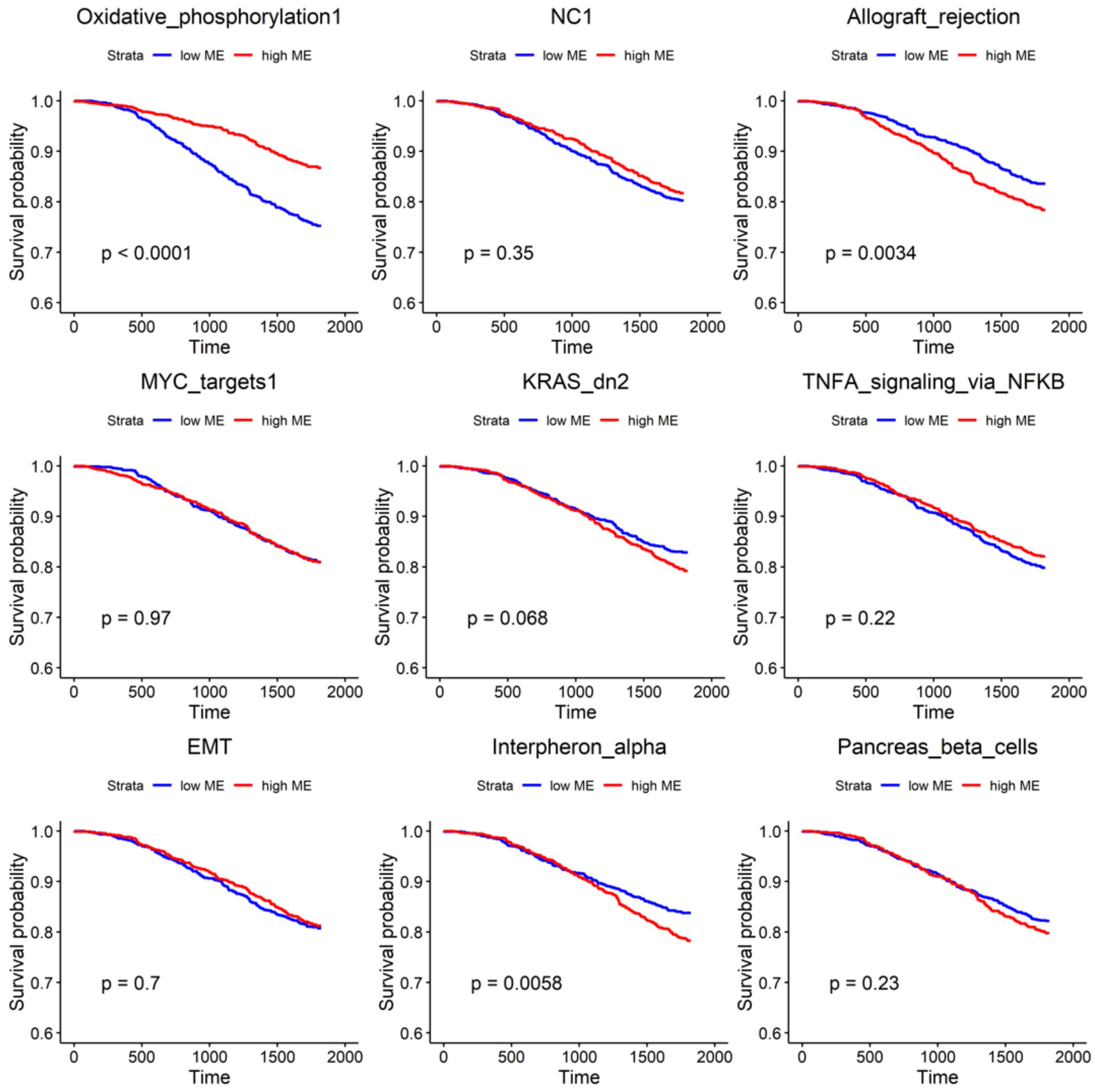


**Supplementary Figure 3.** Kaplan-Meier survival curves for global METABRIC modules (part 1). ME = Module Eigengene.

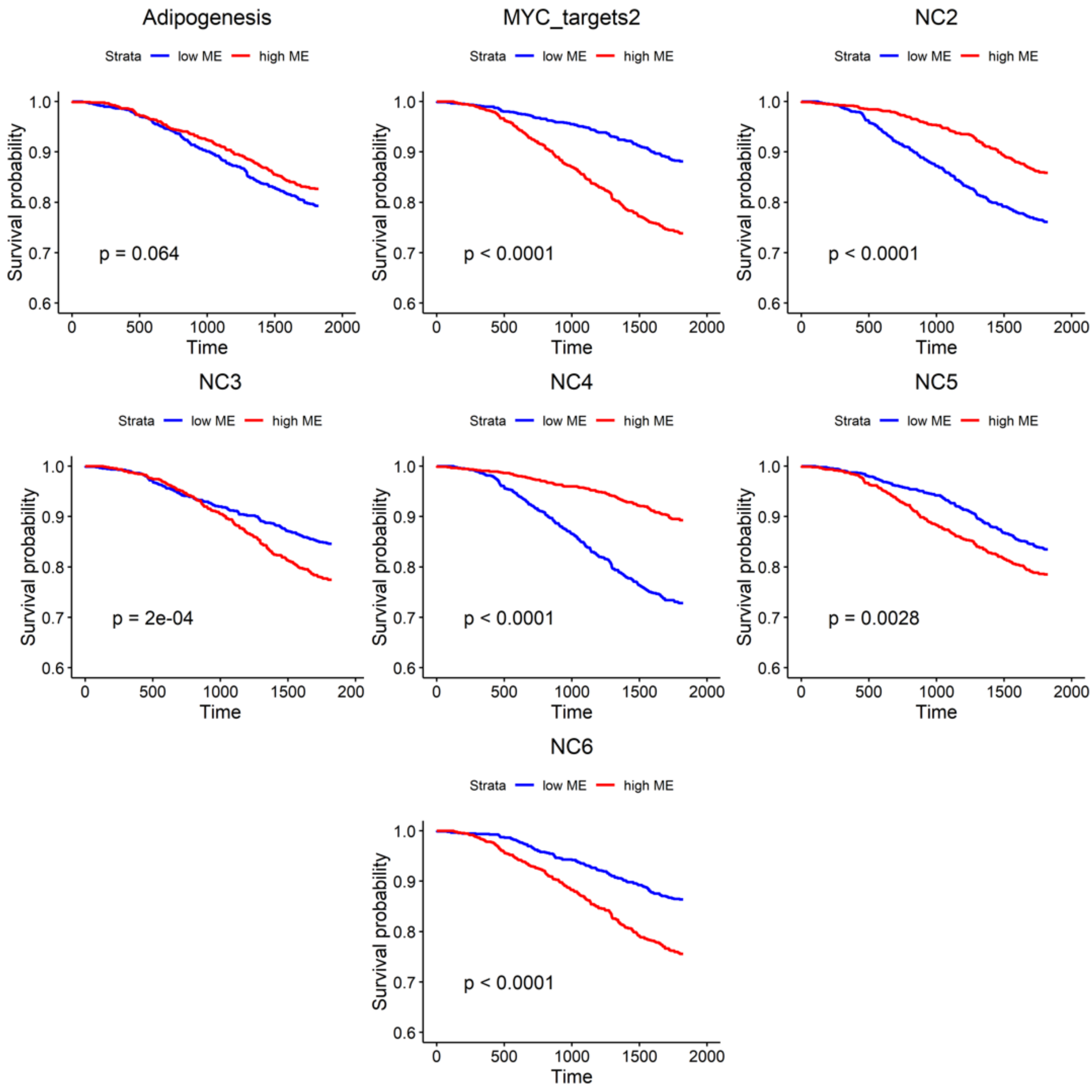


**Supplementary Figure 4.** Kaplan-Meier survival curves for global METABRIC modules (part 2). ME = Module Eigengene.

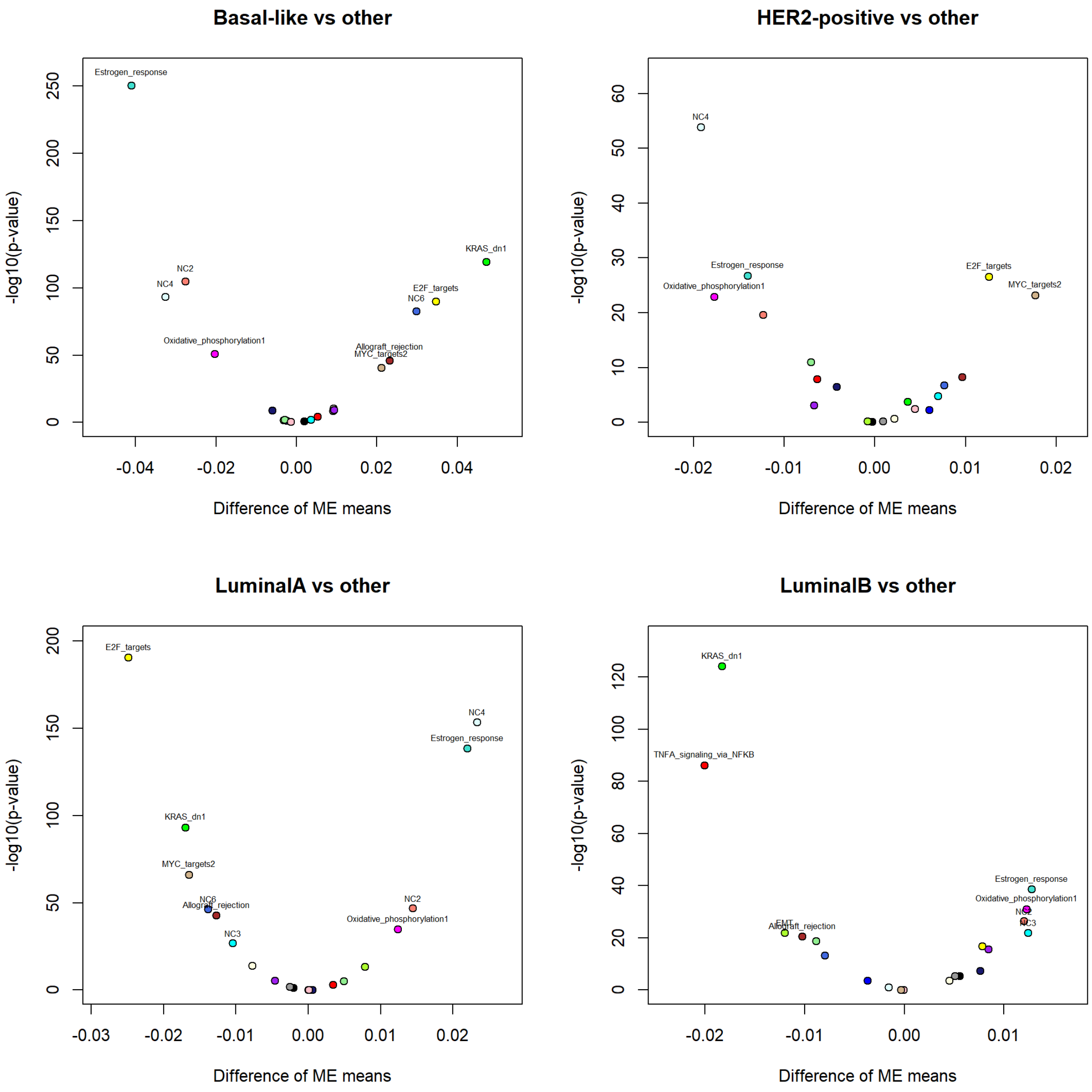


**Supplementary Figure 5.** Difference of modules' expression across subtypes. The expression of each module in a subtype (specified in plot title) is compared with the expression of the same module in all other subtypes pooled together. X-axis shows the difference between module eigenene between the two groups, y-axis represents the significance for the comparison.

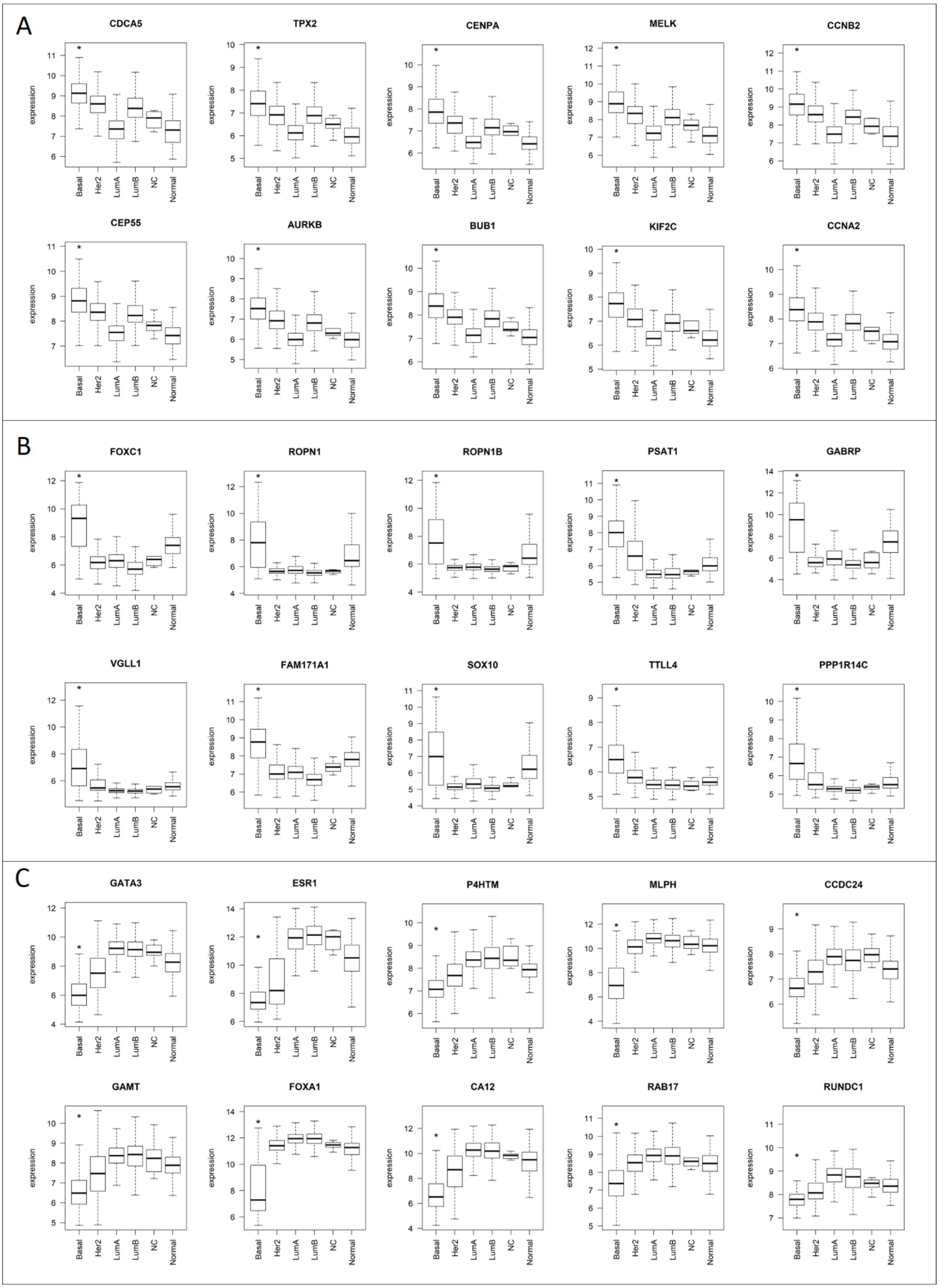


**Supplementary Figure 6.** Expression of the first 10 hubs of E2F_targets (A), KRAS_dn1 (B) and Estrogen_response (C) modules across subtypes. * p-value for the comparison between basal-like and all other subtypes pooled together $<2.2*10^{-16}$

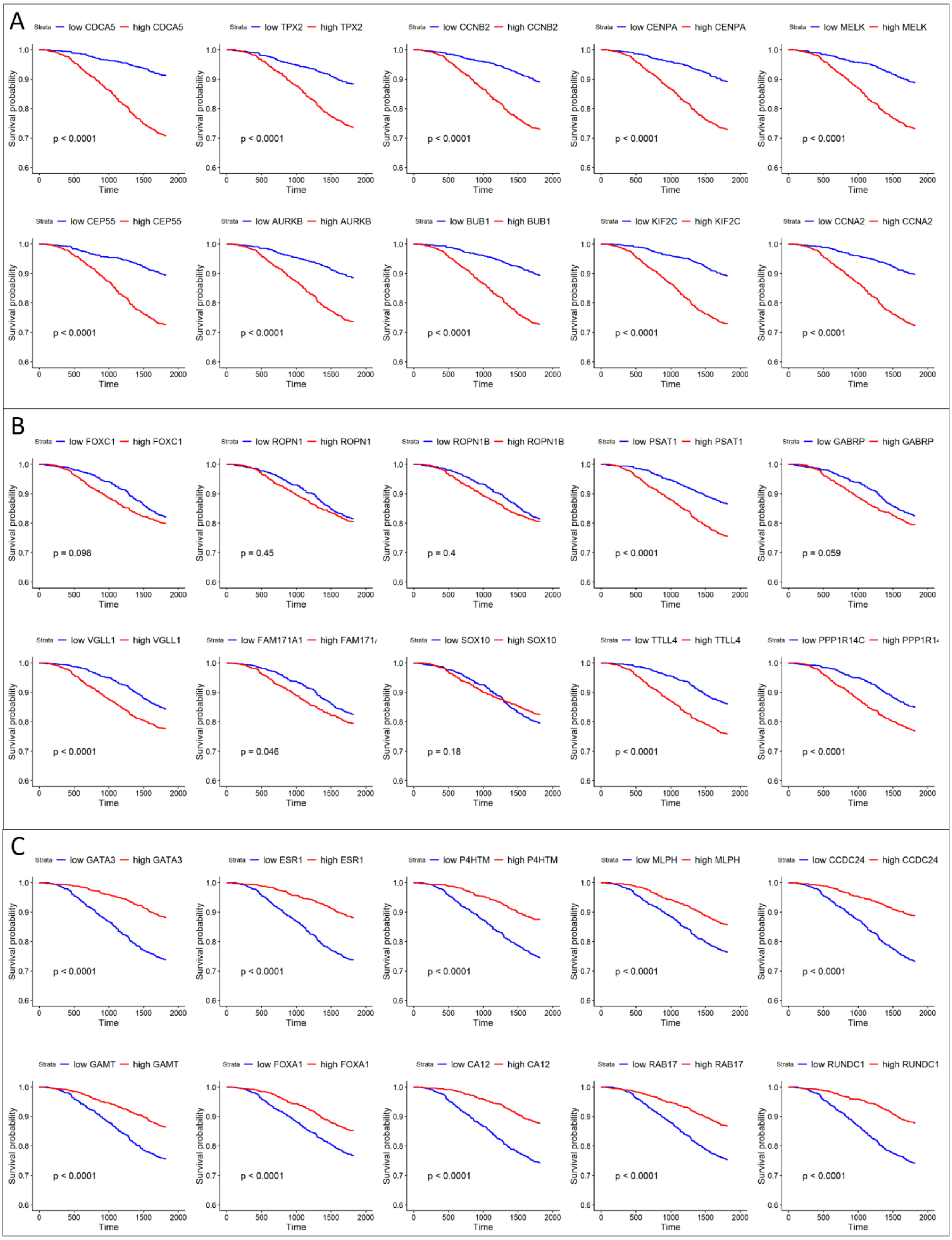


**Supplementary Figure 7**. Kaplan-Meier survival curves indicating the survival probability for patients with either high or low expression of a specific gene, indicated in the legend. First 10 hubs of E2F_targets (A), KRAS_dn1 (B), and Estrogen_response (C) genes.

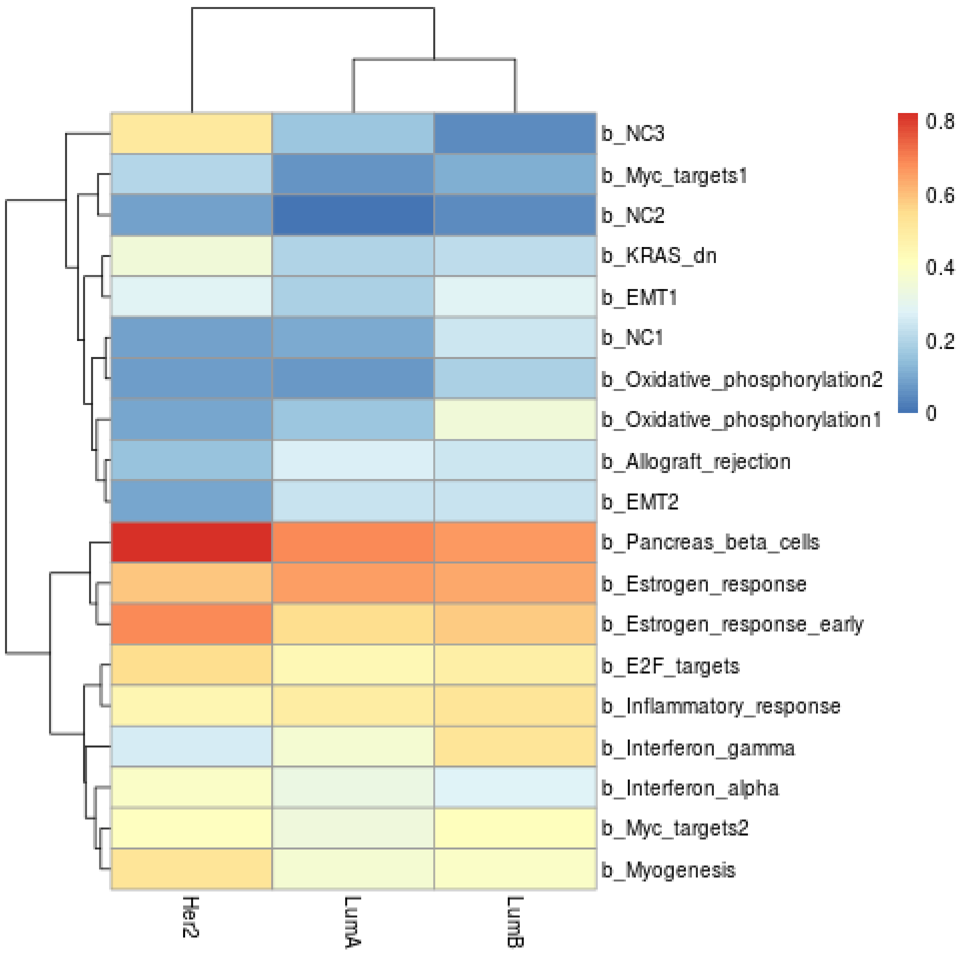


**Supplementary Figure 8.** Percentage of basal modules' genes in the "unconnected" compartment of other subtypes. Red indicates high percentage, blue low percentage. Different modules are in rows, subtypes in columns.
